# Interdigitated dynamic actin protrusions maintain stable cadherin adhesion

**DOI:** 10.1101/2020.10.12.335489

**Authors:** John Xiao He Li, Vivian W. Tang, William M. Brieher

## Abstract

Cadherins build stable, cohesive sheets of cells using paradoxically weak bonds. Actin is thought to convert weak binding into strong adhesion either by transmitting myosin dependent pulling forces to adhesive junctions or by clustering cadherins in the plane of the membrane. Here, however, we show that continuous actin polymerization stabilizes cadherin adhesion by directly driving membrane protrusions, not by promoting contractility or cadherin clustering. Lateral membranes of epithelial cells are continuously pushed against each another by protrusions. Micrometer sized cadherin puncta, long thought to be clusters of cadherins, turn out to be patches of microspikes interlocked by cadherin homophilic bonds to hold neighboring cells together. When actin polymerization is blocked, protrusions cease, puncta disappear, and lateral membranes detach from one another. In contrast, inhibiting myosin II contractility has no effect on adhesion.

**One Sentence Summary:** Stronger together: membrane interdigitations keep cells attached.

## Main Text

Cadherin mediated adhesion requires continuous polymerization of actin cytoskeleton at the adhesion sites, but the reason for this remains a major question in cell biology (*1*). The consensus opinion is that the actin cytoskeleton strengthens adhesion by gathering cadherins into clusters (*2*–*7*) and pulling on them (*8*–*13*); but this contractility model does not account for why adhesion depends on fast polymerization dynamics. Recent results have shown an important role for actin polymerization factors Arp2/3, Ena/VASP proteins, and CRMP-1 in assembling actin at cell-cell junctions in epithelia and endothelia (*13*–*18*). These factors make protrusive actin structures like lamellipodia (*16*). There is then a mismatch in the contractility model between how the junctional actin is made and what it is for (*13*). An alternative model might better explain the relation between actin’s dynamics and function: actin polymerization dependent pushing forces convert weak cadherin bonds into strong adhesion by keeping cadherins from 2 cells close (*19*). Here we test in kidney epithelial cells (MDCK) whether this mechanism can explain the prominent cadherin clusters (often referred to as puncta) that serve as sites for strong cell-cell adhesion (*20, 21*).

E-cadherin puncta appear as “beads on a string” on the lateral plasma membranes between 2 cells in established cell sheets (Fig. 1, B to D) (*22, 23*). Lateral membranes refer to the sides of polarized epithelial cells below the apical junctional complex (AJC) and above the basal plane (Fig. 1A). The lateral domain makes up the vast majority of cell-cell contact area while the AJC is only a narrow strip. The cadherin puncta have membranes twice thicker than the flat regions (Fig. 1E and fig. S1) suggesting membrane enrichment within puncta.

**Figure 1.**
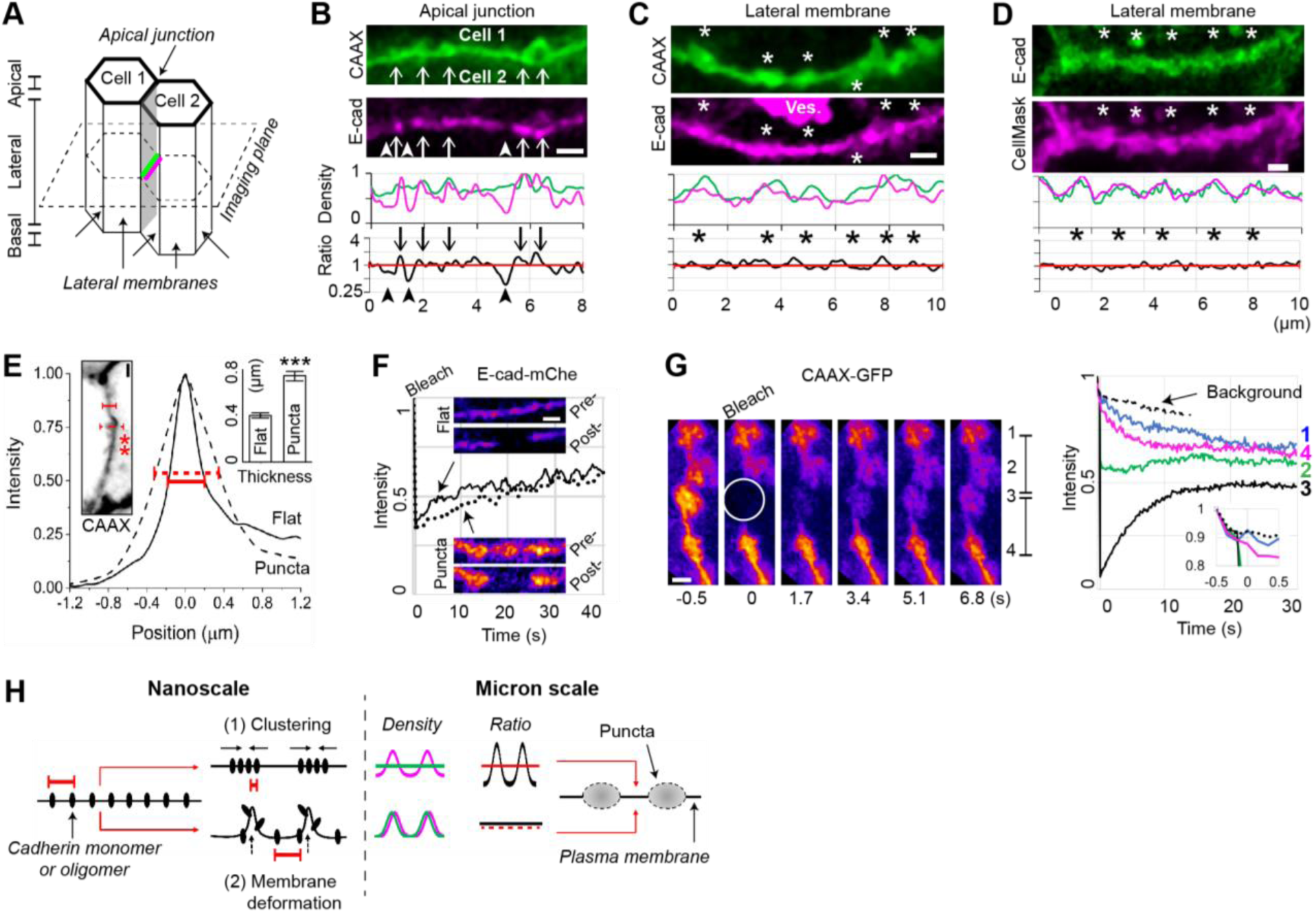
Mesoscale puncta at the cell-cell interface are convoluted plasma membranes. (**A**) 3 domains of epithelial cells. One of the lateral membranes is shaded gray. The horizontal plane indicates the focus plane in **C** to **G**. (**B and C**) Canine kidney epithelial cell sheets co-expressing a plasma membrane-targeted green fluorescent protein (CAAX-GFP) and an E-cadherin fused to mCherry (magenta). Arrows, microspikes at apical junctions (*18*). Arrowheads, voids between E-cadherin puncta. Asterisks, membrane puncta. Ves, intracellular vesicles. The plots show the density of markers along membranes and the cadherin:membrane ratio (red line = 1). (**D**) E-cadherin-GFP (green) is co-stained with a lipophilic dye (magenta). (**E**) The curves show the averaged linescan orthogonal to the membrane at flat regions (solid line) and puncta (dashed line and asterisks). Inset plot, mean membrane thickness (± SEM). *N* = 30 flat regions and 58 puncta in 6 cells. *** *P* < 6×10^−8^, 2-sided t-test. (**F**) Fluorescence recovery after photobleaching (FRAP) of E-cadherin in a flat region (solid line) and a punctum (dashed line). The plot shows the intensity corrected for background bleaching. (**G**) FRAP of the lipidated GFP. Punctum #3 (circle) is bleached. The plot shows the intensity of each punctum (solid lines) and background (dashed line). Inset, −0.5 to 0.5 s. (**H**) Two models to explain mesoscale cadherin puncta (color coding of density and ratio plots same as **B** to **D**). All scale bars, 1 μm.

The widely accepted clustering model to explain the cadherin puncta does not require the enrichment of membranes. Yet, membrane-deforming processes do (*24, 25*). Thus, membrane enrichment in the cadherin puncta can distinguish whether the puncta are due to molecular clustering or membrane deformation (Fig. 1H). 2-color imaging shows that a plasma membrane-targeted green fluorescent protein (GFP) as well as a lipophilic dye co-accumulates with E-cadherin in the puncta (Fig. 1, C and D, and fig. S1). These lipophilic markers do not bind to cadherins or form membrane microdomains (fig. S1); yet they appear as mesoscale puncta like cadherins, which can only be explained by convolution of the membrane itself.

The close-to-1 cadherin:membrane ratio throughout the lateral membrane clearly favors the alternative model that cadherin puncta are actually convoluted membranes (Fig. 1, C and D). At apical junctions, the cadherin:membrane ratio fluctuates more; this ratio is around 1 at where cadherins appear on the finger-like microspikes but as much as 3-fold lower at the voids between cadherin puncta (Fig. 1B). Apical junction cadherin puncta are known to be actin protrusions (*18*) while the voids are transient junctional defects that await to be repaired with cadherins delivered by microspike protrusions (*15, 18*). The voids are not due to depletion by clustering in the nearby puncta because cadherin clustering depends on homophilic bonds (*2, 4*), yet apical junction puncta are insensitive to low calcium treatment that breaks those bonds (fig. S2). Thus, even at apical junctions, cadherin puncta still do not fit the clustering model because cadherins fail to concentrate more than the membrane. Lateral membranes show no obvious defects between cadherins, suggesting lateral junctions may be maintained differently from the repair mechanism at apical junctions.

Another prediction of the clustering model is that clustering makes cadherins more stable and less mobile (*20*). We used fluorescence recovery after photobleaching (FRAP) to compare the mobility of cadherins in the puncta versus on the flat membrane. The fractions of mobile cadherins are not different between the 2 structures (Fig. 1F, fig. S3, and Movie S1).Thus, mechanistically independent of the nanoscale cadherin oligomerization (*26, 27*), the mesoscale cadherin puncta are not supramolecular clusters as long thought to be (Fig. 1H).

To show that these puncta are a part of the plasma membrane rather than docked vesicles, we checked if lipids diffuse through puncta. First, photobleached lipophilic markers diffuse out of a punctum (labeled #3 in Fig. 1G and Movie S2) causing adjacent puncta to dim, which rules out the possibility of vesicles. Further, diffusion within the membrane should follow Fick’s law where it takes more time to travel a larger piece of membrane. We use this travel time to qualitatively compare the amount of membranes in different structures. For puncta #2 and #4: travel time from the source (#3) is indeed longer to get to #4 which is further away than #2. However, for puncta #1 and #4 that are at the same distance from the source, diffusion takes longer to get to #1. The simplest explanation would be the presence of a large membrane reservoir at punctum #2, which must be traversed to get to #1. Thus, these puncta are highly convoluted, yet non-compartmentalized, plasma membranes.

If not by large scale clustering, how do lateral cadherin puncta become strong adhesion sites? Most puncta are blunt and bead-like but can have pointed tips resembling filopodia and microvilli (Fig. 1C). Actin cytoskeleton is a common cause of membrane deformation (*18*). Simultaneous imaging of membrane and actin shows all puncta, whether they are blunt or pointed, strongly overlap with actin (Fig. 2, A and B, and fig. S4). Importantly, time lapse imaging shows that while blunt puncta are stable over the 0.5-h imaging duration (Fig. 2C, figs. S4 and S5, and Movies S3 and S4), dynamic microspikes frequently extend from and retract into the blunt puncta (Fig. 2, C and D, and Movies S3). To test if the microspikes are driven by actin polymerization, we inhibited polymerization with cytochalasin or latrunculin (*18*). Surprisingly, these drugs not only suppress the microspikes but also remove the blunt puncta (Fig. 2E and fig. S6). Cadherin puncta are thought to be associated with a stable actin population (*20*), yet we found them to depend on a dynamic actin population that is replenished by continuous polymerization.

**Figure 2.**
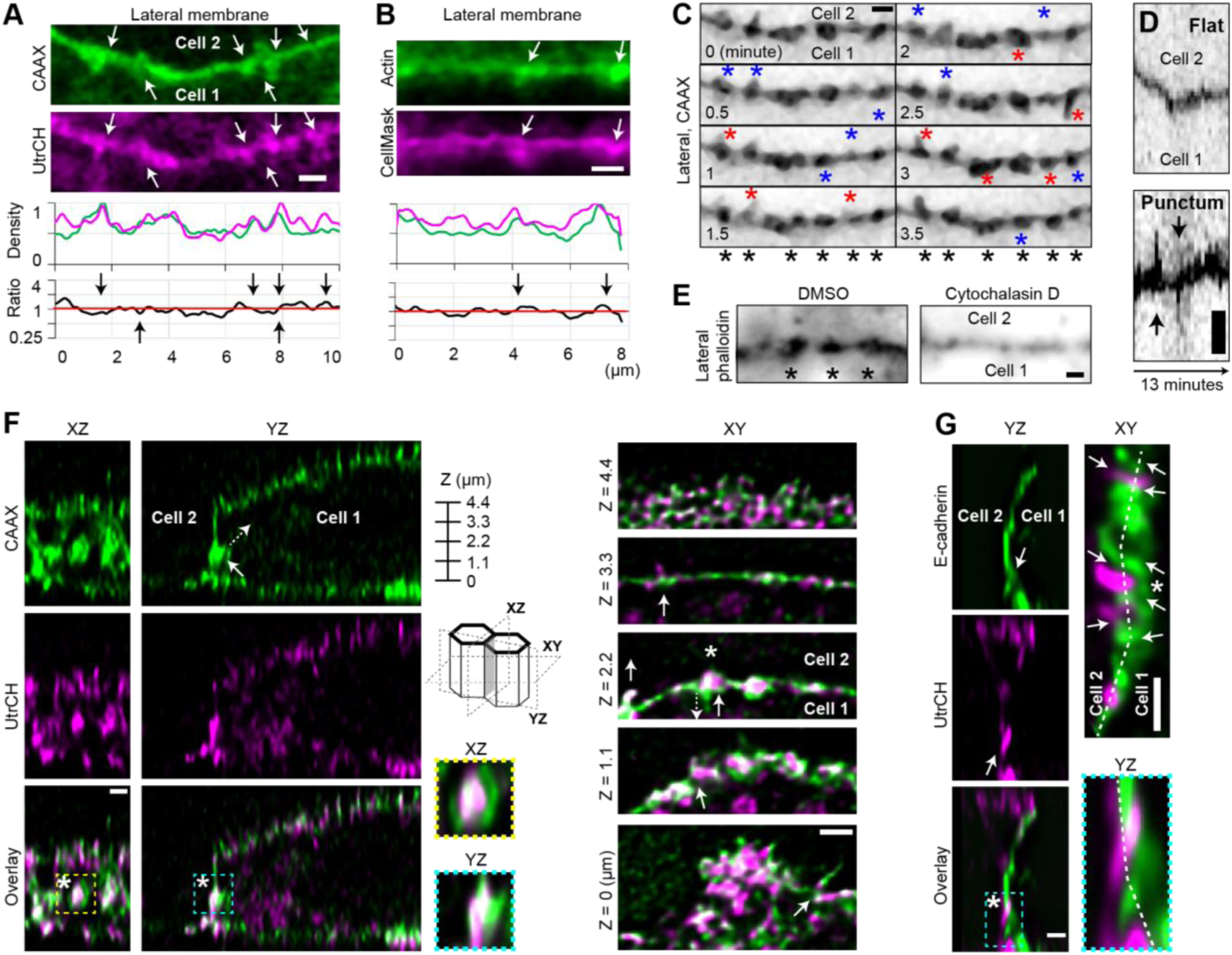
Stable puncta are sites of dynamic actin protrusions. (**A**) Cell sheets co-expressing membrane GFP (green) and the filamentous actin marker, UtrCH (magenta). The plots show the density of markers and the actin:membrane ratio. (**B**) GFP-β-actin (green) and the lipophilic dye (magenta). (**C and D**) Time lapse imaging of membrane puncta. (**C**) All puncta persist throughout the movie (black asterisks); dynamic protrusion (red) and retraction (blue) events occur at these puncta. (**D**) Kymograph drawn perpendicular to the junction shows protrusions (arrows) at puncta but not flat membranes. (**E**) Actin puncta (asterisks) after treated with cytochalasin D (200 nM, 0.5 h) or the vehicle (dimethylsulfoxide). (**F**) Cell 1 co-expresses markers for membrane (green) and actin (magenta) and is next to the non-labeled cell 2. The asterisk shows a punctum that appears as overlapping circles of membrane and actin when viewed en face (XZ); the membrane distends into cell 1 likely to encapsulate a microspike from cell 2 (XY and YZ, arrows). Besides microspikes (arrows), the XY slice at Z = 4.4 µm shows apical surface microvilli; Z = 3.3 µm, apical junction; Z = 2.2 and 1.1 µm, puncta; Z = 0, basal membrane. (**G**) Like **F** but cell 1 is only labeled for membrane (green) and cell 2 only for actin (magenta). Dashed lines, cell boundary. All arrows point in the direction of protrusions. **A** to **E** show XY slices. All scale bars, 1 µm.

The microspikes in Fig. 2C extend towards both sides of the junction, so they might be actin polymerization dependent protrusions or the products of inwardly directed pulling forces. We can distinguish between these two possibilities by labeling the cells on only one side of the junction (*18*). If these structures are due to pushing forces, then the label should only show outwardly directed movement. In contrast, if the structures are due to pulling forces, then we should only see inwardly directed movements. When we coexpressed the membrane and actin markers in cell 1 and left the neighboring cell 2 unlabeled, all the actin microspikes are outside of cell 1’s cell body which is marked by the membrane, not inside of it (Fig. 2F, XY and YZ sections and figs. S4 and S7). Thus, the microspikes are driven by outward protrusive forces of cell 1, not by inward pulling forces. To test if protrusions reach another cell, we mixed cells expressing either of the 2 markers. The vertical section shows that the 2 cells’ lateral membranes are closely touching and their microspikes burrow deep into each other (Fig. 2G and fig. S8). The 2 cells’ microspike protrusions are side by side like interdigitated fingers, thus creating more contact area to form cadherin adhesions.

We wished to understand how microspikes from 2 cells appear in clusters despite their average density across the whole membrane is rather low (fig. S6). Z sections of green/magenta cell pairs show patches of interdigitated microspikes throughout lateral junctions (Fig. 3A). Microspikes are less interdigitated at apical junctions whose “default” state is a continuous belt, so 1 microspike is sufficient for the repair function (Figs. 2F and 3A). In agreement with light microscopy, electron microscopy shows that 95% of membrane protrusions (*N* = 39) engage in a “handshake” configuration (Fig. 3E). Looking at the imaging angle (Fig. 3E, dashed line), one may see as many as 6 layers of membranes around 2 microspikes, which explains the enriched membranes in the puncta shown by light microscopy. Lastly, time lapse imaging of green/magenta cell pairs show that juxtaposed microspikes from 2 cells remain antiparallel to each other, no matter in what directions they are protruding or retracting (Fig. 3D, arrows, and Movie S5). This sustained “handshake” between microspikes means that they are possibly engaged by cadherin homophilic bonds on their tips and shafts (Fig. 3A). In support of this explanation, microspikes disengage when treated with a low calcium medium to break homophilic bonds, even though cells are making similar amount of microspikes (Fig. 3B and figs. S2 and S8). Only 26% of the lateral microspikes in low calcium versus the majority of those under normal conditions (72%) are engaged with another microspike from the neighboring cell (Fig. 3C). On lateral membranes microspikes build cadherin adhesion by becoming interdigitated.

**Figure 3.**
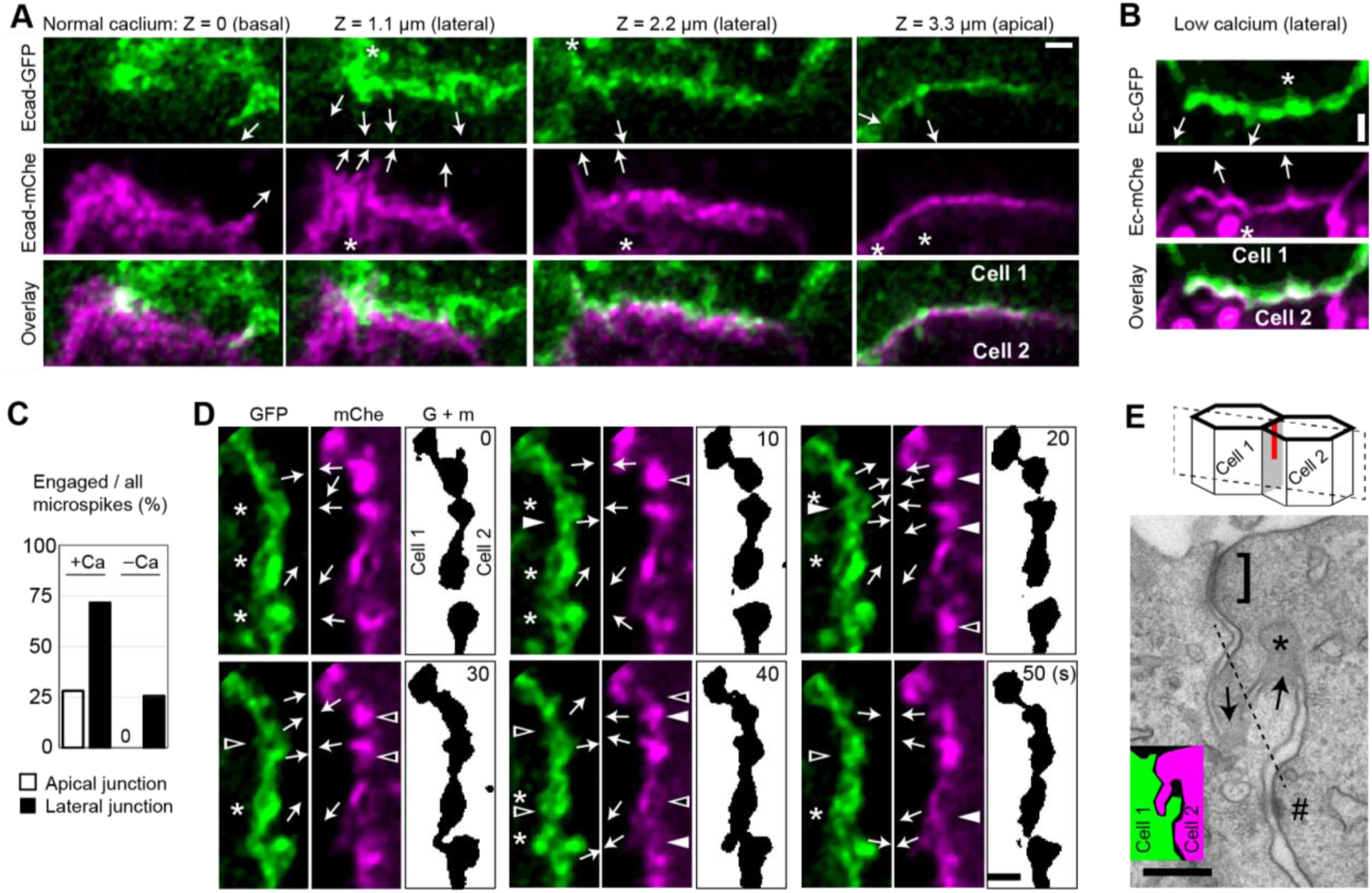
Actin protrusions anchor lateral membranes through interdigitation. (**A and B**) Z sections of 2 cells in a cell sheet expressing the green and magenta E-cadherin markers, respectively. Scale bar, 1 µm. (**A**) 1.8 mM normal media (+Ca). (**B**) 90 µM low calcium media treated for 1 d (–Ca). (**C**) Percentage of microspikes that are engaged with one another. From left, *N* = 32, 106, 14, and 74 microspikes in 2 cells (+Ca) and 4 cells (–Ca). (**D**) Time lapse images of lateral junction between cell 1 expressing E-cadherin-GFP (green) and cell 2 expressing mChe-UtrCH (magenta). The binary image is the sum of 2 colors, which resembles the membrane puncta seen with homogenous labeling (Fig. 2, C and E). Compare the bead-like puncta with the pointed microspikes inside them. Scale bar, 1 µm. (**E**) Vertical thin section electron micrograph of a cell sheet. The image essentially shows the red line in the cartoon. Scale bar, 200 nm. Symbols in all images: arrows, microspike; asterisks, membrane cavities encapsulating the incoming microspikes; bracket, apical junctional complex; #, desmosome; dashed line, a view angle crossing 6 layers of membranes; solid and empty arrowheads, microspike appearance or disappearance, respectively.

The most important implication of microspike interdigitation, however, is perhaps it explains how the dynamic actin cytoskeleton can maintain the stable cadherin puncta. Breaking down the contributions from 2 neighboring cells shows that all the blunt puncta (Fig. 3D, binary images) are patches of pointed microspikes (Fig. 3D, colored images). Puncta look unchanged despite individual microspikes within them appear and disappear all the time; when individual protruding microspikes exceed the length of nearby microspikes, they stick out of the puncta (Figs. 2C and 3D). What we see as cadherin puncta by homogenous labeling are, alas, an optical illusion (Fig. 1).

Actin polymerization inside cells is greatly accelerated by and regulated through various factors. Immunostaining shows the actin polymerization factors Arp2/3, EVL, and CRMP-1 localized in the lateral cortex (figs. S9 and S10). RNA interference and drugs against Arp2/3, EVL or CRMP-1 all decreased the density of microspikes on lateral membranes (Fig. 4, A and B, and fig. S11).

**Figure 4.**
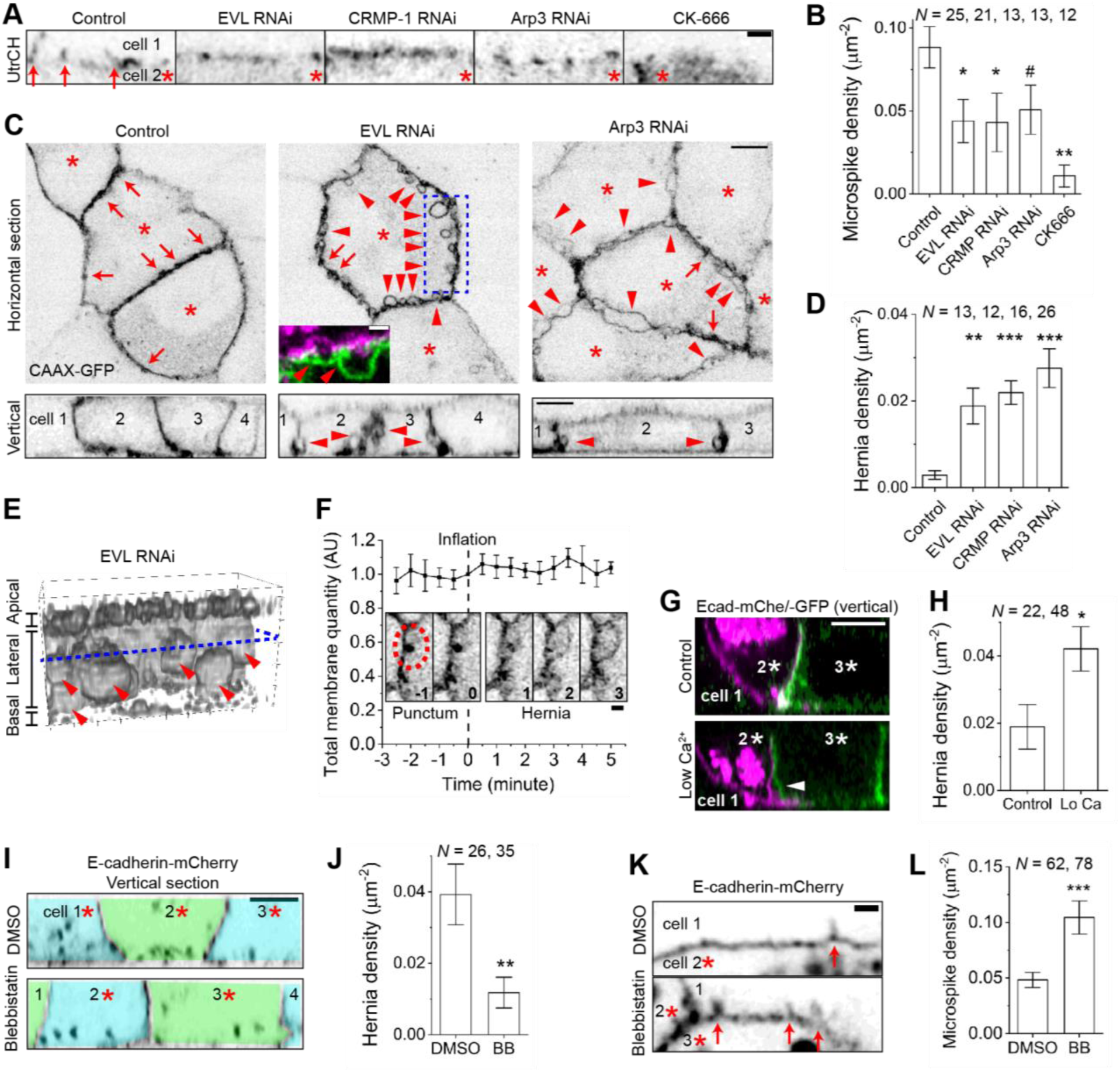
Actin polymerization but not myosin II contractility is required for lateral membrane protrusion and adhesion. (**A to F**) Control, EVL, CRMP-1 or Arp3 knockdown cell sheets, and control treated with an Arp2/3 inhibitor CK-666 (100 µM, 0.5 h). Colored inset, 2 cells labeled green and magenta, respectively (Scale bar, 1 μm). (**B and D**) Density of microspikes (**B**) and hernias (**D**) normalized to the lateral membrane area, pooled from 2 experiments. (**E**) 3-dimensional reconstruction of the blue box in **C**. (**F**) The montage shows a membrane punctum inflating into a hernia in an EVL knockdown cell sheet (oval). The plot shows the conservation of membranes within the oval after the punctum releases the convoluted membranes stored within (mean ± SD, *N* = 9 puncta). (**G and H**) Cell sheets of mixed green or magenta cells treated with normal or low calcium media (90 µM, 1 d). (**I to L**) Cell sheets treated with a myosin II inhibitor (–)-blebbistatin (BB, 50 µM, 0.5 h). All bar plots show mean ± SEM (*N* = number of junctions) with *P* values calculated by 2-sided t-test comparing with control: # *P* = 0.06, *** *P* < 0.001, ** *P* < 0.01, * *P* < 0.05. Symbols in all images: arrows show microspikes in **A** and **K** and membrane puncta in **C**; arrowheads, hernias; asterisks, cells expressing the marker. Scale bars, 1 µm (**A, F, K**) and 5 μm (**C, G, I**).

The consequence of losing microspikes is the severe detachment of 2 cells’ lateral membranes (Fig. 4, C to E, figs. S12 and S13, and Movies S6 to S8). Up to 17% of the total lateral membrane area in the knockdown cells is herniated versus 2% of that in control (see Methods), which can cause spontaneous tissue tearing (*16*). We further showed that membrane detachment is caused by, not just correlated with, decreased microspike density. The evidence are the movies of membrane puncta inflating into hernias (Fig. 4F, fig. S13, and Movie S9). Because puncta are microspikes, herniation happens when the remaining microspikes in the knockdown cells disengage with one another. Conversely, disengaged microspikes in wild type cells can reengage by “search and capture” to keep the membranes attached (fig. S14 and Movie S10). Additionally, disrupting the engagement between microspikes with low calcium media induces membrane detachment (Fig. 4, G and H, and fig. S2). Thus, interdigitated microspikes maintain the cadherin puncta as sites of strong cell-cell adhesion.

Exertion of myosin dependent pulling forces on cadherin clusters is considered a common strategy for strengthening adhesion. Yet, puncta are not clusters of cadherins but clusters of actin microspikes. Nevertheless, we tested whether myosin II is required for lateral junctions. Inhibiting myosin II with blebbistatin does not cause cell detachment in wild type; it further removes hernias (Fig. 4, I and J, and fig. S15). This is because myosin II activity negatively regulates microspikes, shown by the increased microspikes with blebbistatin treatment (Fig. 4, K and L). Thus, myosin II contractility plays no role, or a negative role, to maintain lateral junctions.

Engaged microspikes are long known to the electron microscopists as “cellular interdigitations” (*22, 28*). Our study shows this structure is dynamic and its function is to maintain strong cadherin adhesion. Interdigitations could keep the vast lateral membranes attached in the absence of an actin belt running parallel to the adhesive interface or the myosin II activity; instead, continuous actin polymerization presses the membranes together (fig. S16). Contrary to the previous beliefs (*19, 29, 30*), we show that actin polymerization dependent pushing forces are the primary mechanism of cell-cell adhesion in established epithelia, whereas myosin dependent pulling forces and large scale clustering are dispensable. Importantly, this model offers a direct explanation for why the continuous actin polymerization is essential for stable junctions (*1*).

## Acknowledgments

We thank A. Belmont, J. Chen, and S. Hilgenfeldt for discussion; J. Kemp, A. Nadkarni, and H. Yu-Kemp for reagents; A. Cyphersmith, G. Fried, O. Gholamalamdari, J. Kim, and M. Sivaguru for assistance on imaging; A. Peng for her work during rotation.

## Author contributions

JXHL and WMB conceived the project and wrote the paper; VWT performed electron microscopy; JXHL performed the experiments and analyzed the data.

## Data and materials availability

All data are provided in this article.

## Supplementary Materials

Materials and Methods

Figures S1-S16

Table S1

Movies S1-S10

References (*1-30*)

