## Supplementary material for "Interdigitated dynamic actin protrusions maintain stable cadherin adhesion": Fig. S1-16, Table S1

**This PDF file includes:**

Materials and Methods
Figs. S1 to S16
Table S1
Movie S1. Fluorescence recovery after photobleaching of E-cadherin-mCherry in a flat region (left) and a punctum (right). Frame interval, 0.4 s.
Movie S2. Fluorescence recovery after photobleaching of CAAX-GFP in a punctum. Frame interval, 0.166 s.
Movie S3. Membrane puncta and protrusions labeled by CAAX-GFP. Frame interval is 30 s, shown as min:sec.
Movie S4. The apical junction and a tilted part of lateral membrane in a cell co-expressing E-cadherin-GFP (green) and mCherry-UtrCH (magenta). Neighboring cells are dark. Frame interval is 60 s, shown as min:sec.
Movie S5. The left and right cell expresses E-cadherin-GFP (green) and mCherry-UtrCH (magenta), respectively. Frame interval, 1 s.
Movie S6. 3-dimensional reconstruction of lateral membranes labeled by CAAX-GFP. Control MDCK cell sheet viewed from apical side. Frame interval, 30 s. Compare with Movies S7 and S8.
Movie S7. 3-dimensional reconstruction of lateral membranes labeled by CAAX-GFP. EVL knockdown cell sheet viewed from apical side. Frame interval, 30 s. Compare with Movie S6.
Movie S8. 3-dimensional reconstruction of lateral membranes labeled by CAAX-GFP. EVL knockdown cell sheet viewed from basal side. Frame interval, 30 s. Compare with Movie S6.
Movie S9. A membrane punctum labeled by CAAX-GFP bursts into a hernia. EVL knockdown cell sheet. Frame interval is 30 s, shown as min:sec.
Movie S10. GFP-β-actin (green) and a lipophilic dye CellMask (magenta) showing microspikes on the 2 cells' herniated membranes reengage each other. Frame interval, 1 s.

References (*1-30*)

**Other Supplementary Materials for this manuscript include the following:**

Movies S1 to S10

### Materials and Methods

#### Cell line, maintenance, and transfection

We maintained MDCK II cells (Kai Simons) in MEM media with L-glutamine (Sandra McMaster, Cell Media Facility, UIUC) and 5% fetal bovine serum (Gemini #100-106). The same media plus 25 mM Hepes (pH 7.4) minus Phenol Red is used for live cell imaging. Live cells were plated in glass bottom dishes (Mattek #P35G-0.170-14-C) and immunofluorescence samples on No.1 glass coverslips. We coated all glass surface with 100 μg $ml^{-1}$ Matrigel (Corning #356237).

All knockdown cells are stable cell lines and validated by western blot (fig. S11). Experiments involving UtrCH were transient expression; for other markers stable cell lines were used. To generate stable cell lines, we transfected cells with Lipofectamine 2000 (Life Technologies) and selected with antibiotics and collected all surviving cells as a stable cell line. We plated stable cells for immunofluorescence and live imaging at confluency density (1:2 split from a confluent culture) for 4 days. For transient expression, we split confluent cells 1:4 to get 90% confluency the next day. Then we transfected 1 μg of plasmid DNA per marker without drug selection. We imaged cells 2–3 days post transfection. We only imaged cells with medium fluorescence intensity to ensure good signal to noise ratio and avoid artifacts from overexpression.

#### Plasmids

We used previously validated shRNAs (*1*) against EVL, 5'-GCA GGG ATT CAG CCG GAT AAA; CRMP-1, 5'-GAT GGA TGA GCT AGG AAT AAA; Arp3, 5'-GTA GAT GCC AGA CTG AAA TTA in pLKO.1 puro vector (RNAi Consortium) and used a scrambled shRNA as control for knockdown (Addgene #1864, David Sabatini). Human β-actin was cloned into pLenti-N-terminal GFP vector (*2*). The following plasmids are from Addgene: E-cadherin-GFP (#28009, Jennifer Stow) (*3*); E-cadherin-mCherry (71366, Alpha Yap) (*4*); CIBN-CAAX-GFP (26867, Chandra Tucker) (*5*, *6*); GFP-UtrCH (26737), mCherry-UtrCH (26740, William Bement) (*7*).

#### Drugs

We used cytochalasin D and CK-666 (#SML0006) from Sigma; latrunculin A (#10010630), (–)-blebbistatin (#13013), ML-7 (#11801), and Y-27632 (#10005583) from Cayman with 0.1% (v/v) vehicle DMSO for control. The effects of drugs are validated in figs. S6, S9, S10, and S15.

#### Antibodies

We raised rabbit polyclonal Ab/pAb against human EVL or CRMP-1 full length recombinant proteins (Pacific Immunology) which have been validated previously (*1*). E-cadherin rat monoclonal Ab/mAb (DECMA-1, sc-59778, used at 1 μg $ml^{-1}$ for immunofluorescence/IMF) and Arp3 goat pAb (sc-10132, 1 μg $ml^{-1}$ for western blot/WB) are from Santa Cruz Biotechnology; p34 rabbit pAb (#07227, 1 μg $ml^{-1}$ for IMF/WB) from Millipore; β-actin mouse mAb (66009-1-Ig, 2 μg $ml^{-1}$ for IMF and 0.04 μg $ml^{-1}$ for WB) from Proteintech; AlexaFluor-labeled goat cross-absorbed secondary antibodies from Thermo (1 μg $ml^{-1}$ for IMF). ZO-1 rat mAb (Dan Goodenough, DSHB #R26.4C) and E-cadherin mouse mAb (Barry Gumbiner, DSHB #rr1) was produced as supernatant from hybridoma in house (James Kemp) and used at the saturation concentration for staining.

#### Microscopy

1) Zeiss LSM 880 scanning confocal microscope with Argon/DPSS/HeNe excitation, an Airyscan detector and a 63×, 1.4 NA objective (voxel size is 42.6 × 42.6 × 187 $nm^3$ for 488 nm); 2) wide field microscope (AxioImager. M1, Zeiss) with Colibri illumination, a CCD camera

(ORCA-ER, Hamamatsu) and a 63×, 1.4 NA objective (voxel size is $0.1 \times 0.1 \times 0.3\ \mu m^3$); 3) V3 OMX microscope (Applied Precision) with 488/561 nm laser excitation, a 100×, 1.4 NA objective and 2 EMCCDs (voxel size is $80 \times 80 \times 200\ nm^3$). We used 1 and 2 for immunofluorescence; 1 for all live imaging except for fig. S5 which is done with 3.

**Immunofluorescence**

We followed the protocol of Louise Cramer and Arshad Desai (https://mitchison.hms.harvard.edu/files/mitchisonlab/files/fluorescence_procedures_for_the_actin_and_tubulin_cytoskeleton_in_fixed_cells.pdf). Membrane hernias and p34 antibody staining require methanol fixation (methanol at -20 °C for 3 minutes). After fixation, methanol was quickly diluted with the Tris Buffer (25 mM Tris pH 7.4, 150 mM NaCl) by 2-fold (v/v) to bring methanol concentration down to 50%, 25% and then completely replaced by Tris Buffer. Phalloidin and E-cadherin staining for drug experiments are done by fixation-then-permeabilization unless noted otherwise: we fixed cells with 1% or 4% EM grade formaldehyde (Electron Microscopy Services #15710, 16%) in the pre-warmed Cytoskeleton Buffer (10 mM MES pH 6.1, 138 mM KCl, 3 mM $MgCl_2$, 2 mM EGTA) for 15 minutes and then permeabilized in 0.02% saponin (m/v) in the Tris Buffer. We used 4% formaldehyde for phalloidin only staining; 1% give stronger staining than 4% for E-cadherin and ZO-1 and was hence used for co-staining these markers. For simultaneous fixation/permeabilization we fixed in 1% formaldehyde in pre-warmed Cytoskeleton Buffer with 0.02% saponin and 1 $\mu g\ ml^{-1}$ dark phalloidin for 5 minutes (*8*). Staining and wash are routinely done in Tris Buffer with 0.02% saponin. We used TRITC- (American Peptide) or Alexa 647-phalloidin (Cell Signaling Technology #8940) at 1 $\mu g\ ml^{-1}$. We mounted fixed sample slides in Prolong Gold antifade (Thermo).

**Immunofluorescence quantification**

To quantify apical junction versus lateral junction staining, we used ZO-1 to generate a web-like mask of apical junction. The inverse of this mask is each cell's interior and used as the mask for lateral junction which often tilt towards cell interior in wild field microscopy. We enlarged the cell interior by 0.9 µm and took the created rim as the apical junction region of each cell. (fig. S2) We measured the mean intensity of each cell at apical junction from the maximum projection of 2 z slices at the apical junction level, and the mean intensity of each cell at the lateral junction from the maximum projection of the whole z stack.

**Live cell imaging**

We used the LSM 880 microscope with heating at 37 °C degree and 5% $CO_2$ incubation. For 2-color simultaneous imaging, we used a dual-pass emission filter. We made sure there is no green/red bleed-through; otherwise sequential acquisition with single-pass filter is used. We used the automatic setting for 3D Airyscan processing. To reconstruct the en face view of puncta on the lateral membranes (vertical sections), we deconvolved the z stacks with the Meinel method (Zen Blue 3.0, Zeiss). For long term imaging, we took care to avoid photo damage: 30 s interval for 5 Z slices/20–60 frames. Z stacks should span to cover the apical and basal membranes: MDCK cell sheets physiologically contract and fluctuate along Z-axis on the cycle of 5–10 minutes which can cause small structures like puncta to "disappear" while it is only moving out of focus. The images shown are taken from the frames free from such cell height fluctuation.

For fig. S4, we used the OMX microscope with heating at 37 °C degree and 5% $CO_2$ incubation. The 2 colors are acquired simultaneously. At 1-minute intervals, we took 3 z slices 200 nm apart for deconvolution but only used the middle slice for analysis. We deconvolved images with SoftWoRx software (Applied Precision) using the Additive method with automatic iteration settings.

**Membrane density plots**
We labeled the lipids with the CAAX motif of Ras fused to GFP (farnesylated) by transfection or the CellMask Orange plasma membrane dye (Thermo) by adding to the cells immediately before imaging. Previously, myristoylated (*9*) and palmitoylated (*10*) GFP probes also showed mesoscale puncta. We drew intensity linescan (line width set to 0.5–0.8 µm cover the thickness of puncta/protrusions) along the membrane to show the line density of the membrane. We normalized linescan by setting [blank background intensity, linescan maximum pixel value] to [0, 1]. The pixel-wise ratio of cadherin:membrane or actin: membrane is calculated by ratio = $(I_1/I_2)/(\mu_1/\mu_2)$, which is dividing their density ($I$) pixel by pixel and normalizing to the ratio of the mean of each linescan ($\mu$). By that we set the mean of the ratio to ~1 while preserving large deviations like the voids at apical junction.
**Membrane thickness**
We measured the membrane thickness at the border between marker expressing and non-expressing cells because the cytoplasmic intensity is not always zero and cannot be used as the background value (*11*). The labeled cell is the right part of the curve thus explaining the higher background. We drew by hand 3-pixel wide lines orthogonally to the membrane at the flat and the center of punctum regions in ImageJ and place the midpoint on the membrane midline. We normalized the intensity linescan curve by setting [the mean of the left most 10 pixels, maximum] to [0, 1] and registered its maximum to position = 0. The maximum search is limited within 10 pixel on either side of the linescan midpoint. We then search for the half maximum of the left half of the curve and fitted its 2 closest data point to a straight line; the intersection of this line with the actual curve gives an approximately half width at half maximum. The plotted membrane thickness is twice of the half width.
**Fluorescence recovery after photobleaching**
We did FRAP of CAAX-GFP in control cells. We did FRAP of E-cadherin in EVL RNAi cells to be able to find hernias. We bleached a circle of 1 µm in radius to visually dark which is 10% and 25% of the initial intensity with 405 nm and 488 nm laser for CAAX-GFP and E-cadherin-mChe, respectively. The frame interval is 0.166 s for CAAX and 0.4 s for E-cadherin. The intensity curve is not normalized to background for CAAX because we wanted to show the difference between each region of interest (ROI) and the background due to receiving dark molecules from the bleached region; each ROI is normalized to its own initial intensity. We normalized the intensity of E-cadherin in the ROI to the background (the entire image minus the twice enlarged ROI) to correct for background bleaching; we then fitted the recovery curve to a single exponential process of time ($t$) in Origin (*12*), $Intensity = A(1 - exp(-(t-to)/\tau))$. Half time = $0.7\tau$; mobile fraction = $A$. We designed a way to compare the membrane quantity in the ROIs given the unequal expression level of the marker across cells: we normalized the sum intensity of the circular ROI (fig. S3, “Bleaching”) to another circle of unbleached, flat membrane region (“Reference”) as the latter is proportional to the expression level. This allows comparison of membrane quantity in puncta of different apparent brightness (fig. S3B). We used hernias as a true free surface for comparison.
**Low calcium treatment**
Low calcium media (90 µM $Ca^{2+}$) is made by adding FBS (1.8 mM $Ca^{2+}$) to 5% final concentration in calcium-free MEM (Sigma #M4767). This concentration severely disrupts trans interactions between cadherins on 2 cells (*14–17*) without affecting the intracellular calcium concentration (*18*). In addition to immunostaining, its effect was confirmed by a tight junction permeability assay. 6-d confluent cell sheets on coverslips pre-treated with normal or low

calcium media for 1 d were incubated with an anti-E-cadherin antibody (rr1) in the respective media for 1 h. After the media was aspirated, cells were fixed by 1% formaldehyde in the Cytoskeleton Buffer for 15 minutes and quenched in the Tris Buffer. Importantly, cells were not permeabilized with detergent but incubated directly with Alexa 647 conjugated secondary antibody against mouse IgG for 1 h (avoid green dyes because of the tissue autofluorescence). Coverslips were then mounted in the Tris Buffer and imaged immediately for the 647 nm (signal) and phase channels (tissue integrity). Cells sheets in the normal media without rr1 antibody were used as a negative control and its staining intensity (background) is set to zero (fig. S8D).

**Microspike engagement quantification**

Lateral microspikes often appear in more than 1 z slice due to limited axial resolution, which leads to overcounting and thus prevents automatic microspike detection used for apical junction (*9*). We manually detected microspikes after blinding the images' filenames. Microspikes extending from the cortex by > 0.5 µm are counted. We assigned the center of each microspike as its position in ImageJ; then the distance between each green/magenta pair of microspikes was calculated by a Macro and filtered by a threshold. This threshold is set to 0.5 µm (twice of the radius of microspikes at the base, ~0.25 µm) meaning only microspikes within 0.5 µm of one another are considered "clustered". We then calculated the ratio between engaged to all microspikes.

**Microspike density quantification**

For the reasons mentioned in "clustering quantification", we manually counted microspikes after blinding the images' filenames. The key is to only count cells with medium UtrCH expression level to avoid artifacts. The expression level of membrane markers does not affect microspike density, but dim images can lead to undercounting. We assigned each microspike its position in ImageJ; then their z positions were recorded by a Macro and assigned as apical or lateral based on the position of the apical junction. The slices 1 µm below the apical junction are counted as lateral. We then calculate the area of lateral membranes which are often tilted and thus deviate from the cylindrical approximation (cell perimeter × height) by ~40%. Instead, we manually drew segmented lines on the membrane contour at the z positions where tilting starts or turns. Then the lateral area between 2 z slices was calculated by a trapezoidal approximation: $area = \text{mean length of 2 segmented lines} \times \sqrt{(\text{height difference}^2 + \text{distance between centers of mass of 2 lines}^2)}$. The total area is the sum of these trapezoids. But very tilted, close-to-flat lateral membranes should be avoided because microspikes do not stick out in these regions, leading to undercounting. For this reason, we also did not count microspikes on the basal cryptic lamellipodia. Finally, we normalized the number of microspikes by the membrane area. The maximum packing density of microspikes (4.8 $\mu m^{-2}$) is estimated by considering microspike cross sections as circles of 0.5 µm in diameter.

**Hernia quantification**

For the same reason as for microspikes, we manually counted hernias after blinding the images' filenames. Hernias are counted from z stacks, not vertical sections. Membrane distention with visible hollow cavity is counted as hernias. We normalized the number of hernias by the membrane area to give the density. To estimate the fraction of lateral membrane covered by hernias by hernia density, we simplified hernias as half domes and thus $Fraction = 2\pi r^2\ Density$, where $r$ is the hernia depth (~1 µm).

**Electron microscopy**

MDCK cells grown on Transwell-Clear were processed for EM (*19*, *20*). Briefly, transwells were chilled at 4°C for 6 h before fixation with 3.75% glutaraldehyde, 150 mM NaCl, and 20 mM Hepes, pH 7.5, at 4°C for 18 h, and then quenched with 50 mM glycine and 150 mM Hepes, pH 7.5, on ice for 1 h. Transwells were rinsed in ice-cold distilled water three times, secondary fixed with 1% osmium tetroxide/1.5% potassium ferrocyanide for 2 h on ice, rinsed four times in ice-cold distilled water, en bloc stained with freshly prepared and filtered 2% uranyl acetate in distilled water on ice for 2 h, and rinsed four times in ice-cold distilled water. Transwells were dehydrated with sequential 5-min incubations in 50, 75, 95, 100, 100, and 100% ethanol at room temperature. Epon-Araldite (EMbed 812) was added to transwells and allowed to polymerize at 60°C for 48 h. Ultrathin sections were cut using a microtome (UltraCut S; Reichert), layered onto carbon-coated copper grids, and stained with freshly made/filtered 2% lead citrate. Images were collected with a microscope (1200EX; JOEL, Ltd.) at 60 kV. The protrusions are counted as clustered only if they are immediately juxtaposed.

**Statistical test**

2-sided t-test was used to compare a group with control (null hypothesis: the means are equal).

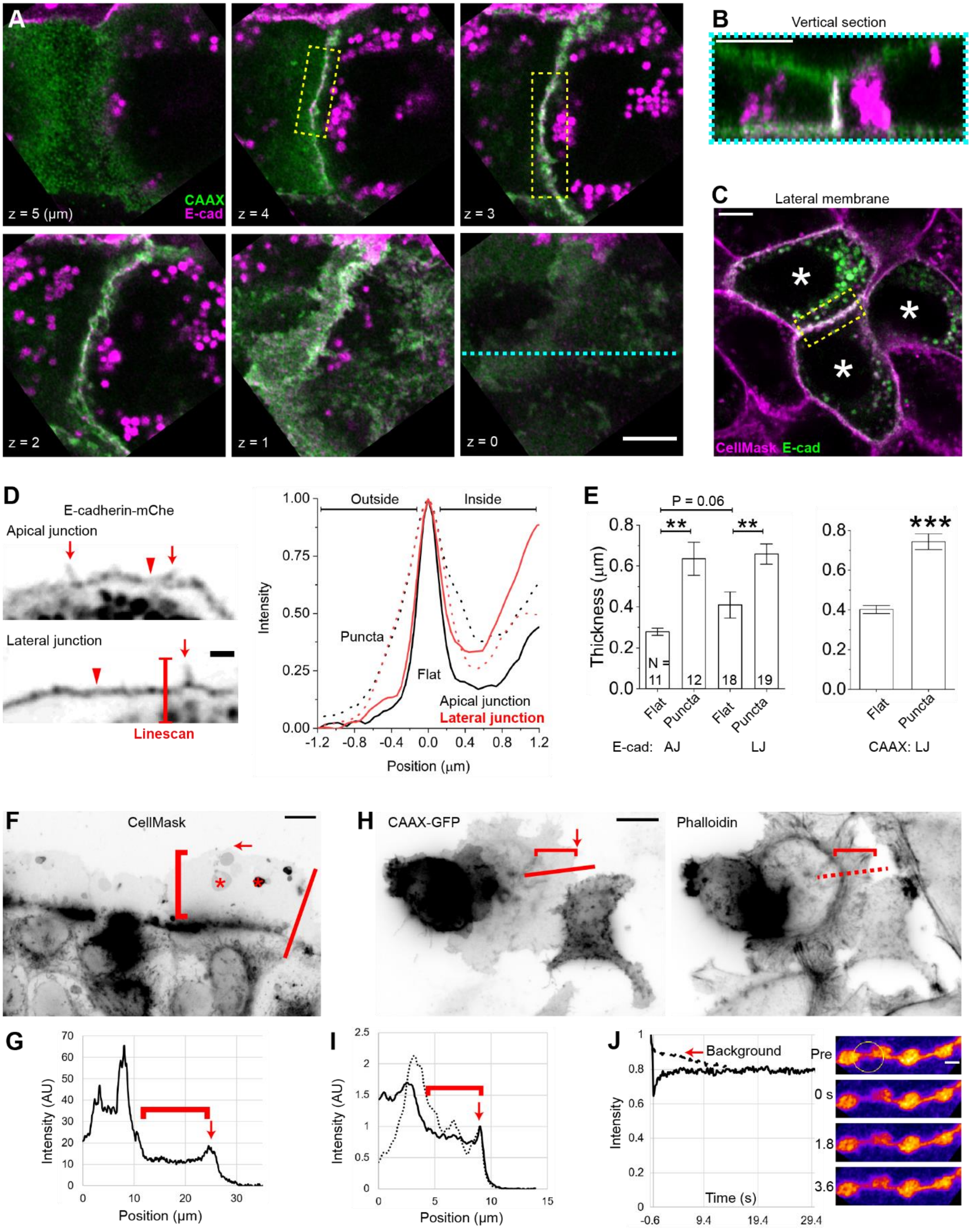

A
CAAX
E-cad
z = 5 (µm)
z = 4
z = 3
z = 2
z = 1
z = 0
B
Vertical section
C
Lateral membrane
CellMask E-cad
D
E-cadherin-mChe
Apical junction
Lateral junction
Linescan
Outside
Inside
Puncta
Flat
Apical junction
Lateral junction
Intensity
Position (µm)
E
P = 0.06
Thickness (µm)
N =
Flat
Puncta
E-cad: AJ
LJ
CAAX: LJ
F
CellMask
G
Intensity (AU)
Position (µm)
H
CAAX-GFP
Phalloidin
I
Intensity (AU)
Position (µm)
J
Background
Intensity
Time (s)
Pre
0 s
1.8
3.6


**Fig. S1**

**Fig. S1. The plasma membrane markers.**

(**A and B**) Canine kidney epithelial cell sheet coexpressing CAAX-GFP (green) and E-cadherin-mCherry (magenta). Z = 4 μm, apical junction; 3 and 2 μm, lateral junction; 1 μm, basal lamellipodia; 0 μm, basal plane. The yellow boxes are shown in Fig. 1. Note that E-cadherin colocalizes with the lipidated GFP everywhere on basolateral plasma membrane but not the apical surface and microvilli, as expected for E-cadherin. (**B**) The vertical section (along the cyan line in **A**) shows lateral membranes of 2 cells are closely touching. (**C**) Live canine kidney epithelial cell sheets sporadically expressing E-cadherin-GFP (asterisks) are stained with a lipophilic dye, CellMask. Yellow box, the junction between 2 green cells is shown in Fig. 1. (**D**) The curves show the average of intensity linescan orthogonal to the membrane at the flat regions (arrowheads and solid lines) and puncta (arrows and dashed lines). (**E**) Mean membrane thickness (± SEM) measured with different markers. The number of linescan ($N$). *** $P < 6\times10^{-8}$, ** $P < 0.01$, 2-sided t-test. Note that the thickness of flat membranes is larger than the theoretical resolution limit of the microscope, $d$ = 125 nm (calculated by $d = 0.61\lambda/1.7NA$ with $\lambda$ = 488 nm, Numerical Aperture = 1.4, and 1.7-fold improvement by the AiryScan detector) but consistent between the CAAX-GFP ($\lambda$ = 488 nm, Fig. 1) and E-cadherin-mChe markers ($\lambda$ = 561 nm), which can only be explained by membrane deformation but not planar clustering. (**F to I**) CAAX-GFP (**F and G**) and CellMask (**H and I**) labeling of truly flat membranes in the lamellipodia. Note the lack of mesoscale puncta or membrane domains. The plots show intensity linescan of the red lines in the images; solid line is membrane and dashed line is actin; the curves' right end is outside the cell. Arrow, membrane ruffling at the leading edge. Bracket, lamellipodia. Asterisks, blebs due to fixation. Cells lively labeled with CellMask are fixed without permeabilization. (**J**) CAAX-GFP is photobleached (circle) to a less extent than Fig. 1 thus recovers to the equilibrium faster with the adjacent membranes (solid curve). Dashed curve shows background bleaching due to imaging. The 2 curves meet means this marker is close to completely mobile thus unlikely associated with any stable membrane microdomains. Scale bar, 5 μm (A to C), 1 μm (D and J), and 10 μm (F and H).

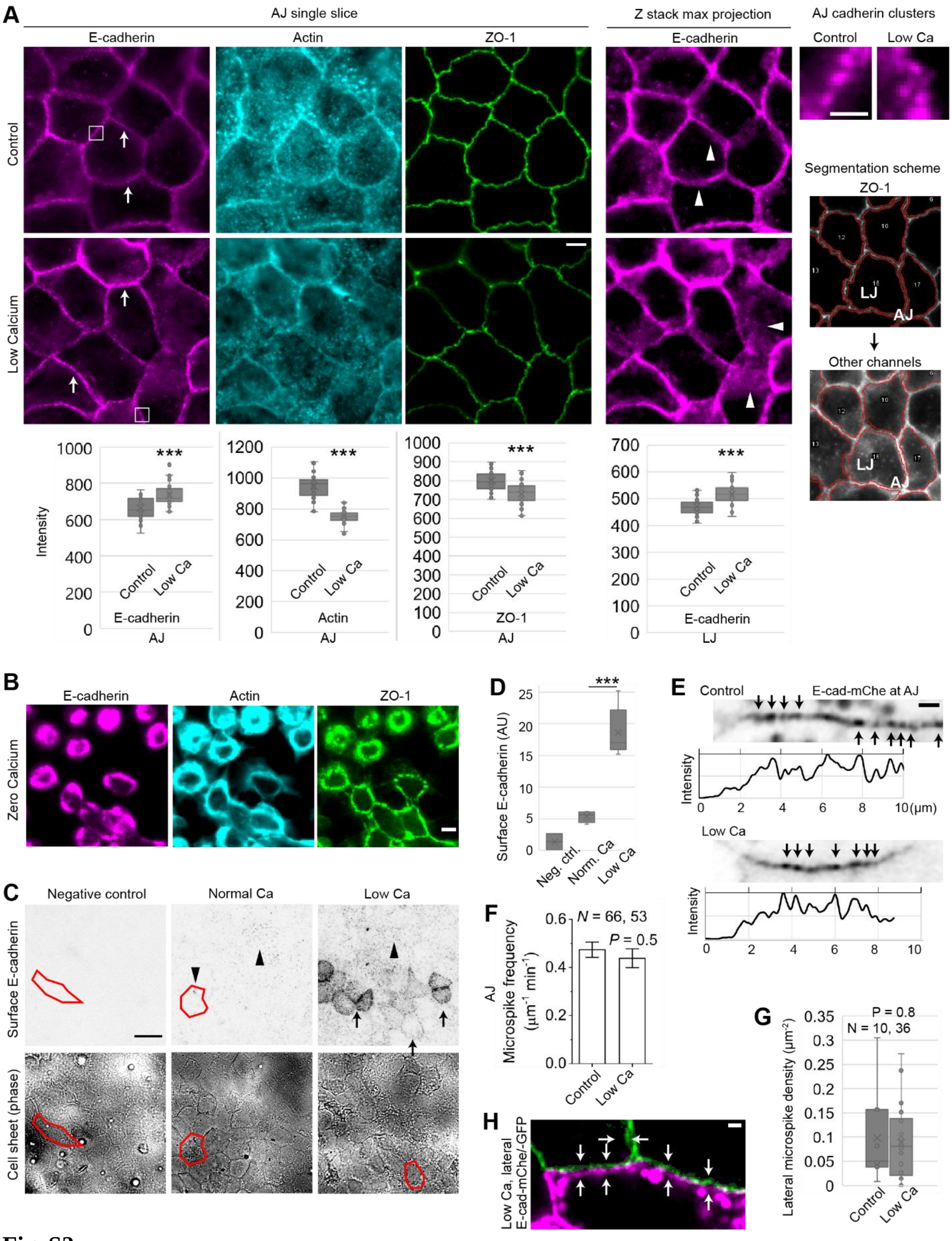
A
AJ single slice
Z stack max projection
AJ cadherin clusters
E-cadherin
Actin
ZO-1
E-cadherin
Control
Low Ca
Control
Low Calcium
Segmentation scheme
ZO-1
LJ
AJ
Other channels
Intensity
***
E-cadherin
AJ
Actin
AJ
ZO-1
AJ
E-cadherin
LJ
B
E-cadherin
Actin
ZO-1
Zero Calcium
C
Negative control
Normal Ca
Low Ca
Surface E-cadherin
Cell sheet (phase)
D
Surface E-cadherin (AU)
Neg. ctrl.
Norm. Ca
Low Ca
E
Control
E-cad-mChe at AJ
Intensity
10 (μm)
Low Ca
F
N = 66, 53
P = 0.5
AJ Microspike frequency (μm$^{-1}$ min$^{-1}$)
Control
Low Ca
G
P = 0.8
N = 10, 36
Lateral microspike density (μm$^{-2}$)
H
Low Ca, lateral E-cad-mChe/-GFP

**Fig. S2**

**Fig. S2. Cadherin homophilic bonds are required for maintaining apical junctions and lateral junctions but not cadherin nanoclusters.**

(**A, C, D**) Validation of low calcium treatment (90 µM, 1 d, which severely disrupts trans interactions (*14–17*)). (**A**) Staining of junctional and cortical actin, and apical junctional marker ZO-1 decrease due to low calcium. Cadherin antibody (rr1) staining increases at both apical junction (arrow) and lateral junction (arrowheads) though adhesive activity is weakened at both junctions shown by the broken tight junction (see **C and D**) and lateral membrane hernias (Fig. 4). Further, the maintenance of both apical junction and lateral junction is actin dependent (*20*); actin is found to be disrupted at both the apical junction and the cortex. Cadherin nanoclusters are not affected (see enlarged images on the far right, scale bar 1 µm), but they are still insufficient for junction maintenance (*16*). Scale bar, 5 µm. The plots show interquartile range (IQR, box) with mean (cross). $N \geq 74$ cells. *** $P < 10^{-4}$, 2-sided t-test. The segmentation scheme shows how apical junction and lateral junction E-cadherin is measured. (**B**) For comparison, zero calcium treatment (phosphate buffered saline plus 3 mM $Mg^{2+}$, 0.5 h) completely dissolves the cell-cell adhesion thus prevents further analysis. Scale bar, 5 µm. (**C**) An assay of cadherin dependent apical junction complex maintenance (*21*). An anti-E-cadherin antibody (rr1) gains access to the lateral surface due to low calcium. The cells grown on coverslips were not permeabilized so that the antibody only stains the surface cadherins. This antibody sparsely binds to the apical surface cadherins (arrowheads) in the normal media but has access to the lateral surface cadherins (arrows) in low calcium. Red polygons trace the shape of 1 cell in the cell sheet. The negative control with only secondary antibody but no rr1 antibody shows no cadherin staining meaning that the staining is specific. Scale bar, 25 µm. (**D**) Surface E-cadherin staining intensity. IQR with mean. $N = 2$ negative control, 4 normal calcium, and 6 low calcium images. *** $P < 10^{-3}$, 2-sided t-test. (**E**) Low calcium has little effect on the E-cadherin-mChe puncta (arrows) at apical junction. Scale bar, 1 µm. The plots show intensity linescan normalized to its own maximum value; the left most part of the curve shows the non-transfected neighboring cells and reads zero intensity. Note the similar amplitude and width of peaks in the 2 plots. (**F**) Low calcium does not change the frequency of microspikes at apical junction. The plot shows mean ± SEM; the number of junctions (*N*) from 2 experiments; 2-sided t-test. (**G**) Low calcium does not change the density of lateral microspikes. The plot shows IQR and mean; the number of microspikes (*N*); 2-sided t-test. This density is ~50 times lower than the density needed to cover the entire lateral surface. At the same density compared to normal conditions, the microspikes still fail to appear next to one another in low calcium. Thus, the interdigitation between microspikes are not entirely due to random localization but specifically enhanced by cadherin binding (Fig. 3). (**H**) Hernias (arrows) due to low calcium are viewed in a horizontal slice. Scale bar, 1 µm.

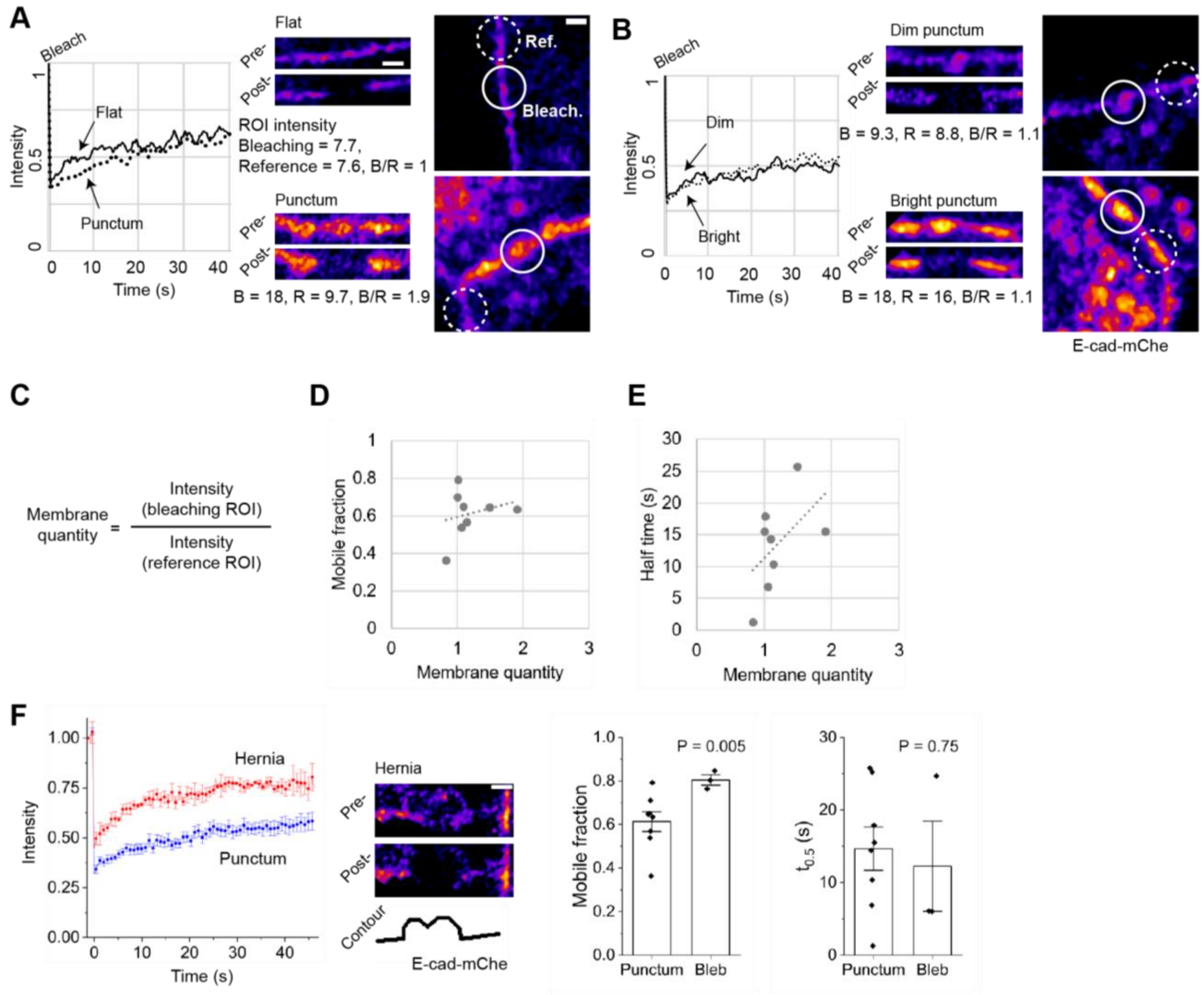


**Fig. S3. Cadherins are not more stable in the puncta than on the flat membranes.**

(**A and B**) Fluorescence recovery after photobleaching of E-cadherin-mChe. The plots show fluorescence recovery after photobleaching corrected for background bleaching. The circles show the ROIs (regions of interest) used to measure membrane quantity by the equation in **C**. The “Reference” ROI (dashed line circle) is an unbleached, flat membrane region of the same size as the “Bleached” ROI (solid line circle). See Methods. (**A**) A flat region and a punctum. Note their mobile fractions are similar (flat, 0.70; punctum, 0.63). Images are used in Fig. 1. (**B**) A dim punctum and a bright punctum. Note their mobile fractions are similar (dim, 0.54; bright, 0.57). (**C**) This equation normalizes the different expression level of cadherins in each cell. (**D**) The mobile fraction is not inversely correlated with either ROI sum intensity before normalization (data not shown) or membrane quantity, contrary to the previous report (*13*). Dashed line, linear fit (Pearson’s $r = 0.25$, $N = 8$). (**E**) The correlation between the membrane quantity and half-life (*22*). Dashed line, linear fit (Pearson’s $r = 0.52$). (**F**) Cadherins on the detached hernia surface are less stable shown by the mobile fraction. The curves show mean ± SEM. $N = 8$ puncta and 4 hernias. $P$ values are by t-test. All scale bar, 1 µm. See Movie S1.

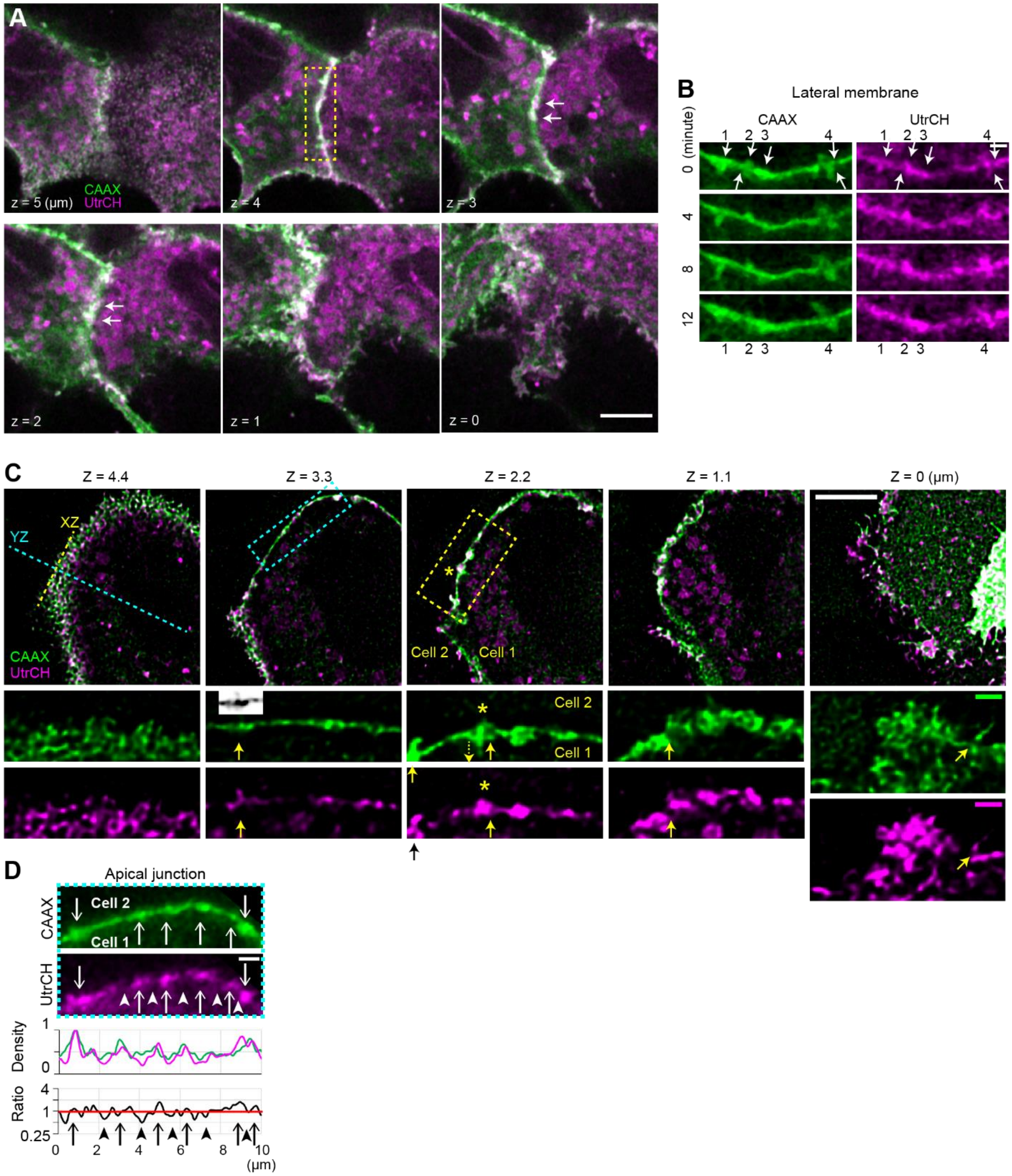


**Fig. S4. Membrane puncta overlap with actin.**

(**A and B**) Lateral puncta (arrows) between 2 cells coexpressing CAAX-GFP (green) and mCherry-UtrCH (magenta, F-actin marker) in a cell sheet. The junction of interest is of homogenous labeling. Z = 5 μm, apical junction; 4 to 2 μm, lateral junction; 1 and 0 μm, basal plane. The yellow boxes are shown in Fig. 2 and panel **B** here. Note that actin colocalizes with the lipidated GFP on both apical and basolateral plasma membranes as expected. (**B**) Time lapse imaging of membrane and actin puncta numbered 1–6. Arrows show their protrusion direction.

All 4 puncta persisted throughout 12 minutes; protrusions shrunk in #1 and 2 and distended in #4. The puncta have much longer lifetime than individual microspikes that last for seconds (Fig. 3). (**C and D**) Cell 1 coexpresses CAAX-GFP (green) and mCherry-UtrCH (magenta). Cell 1 is surrounded by unlabeled neighbors. (**C**) Deconvoluted images. Z = 3.3 μm, apical junction (black/white inset: the dim membrane labeling of the microspike is revealed after adjusting the contrast); 2.2 to 1.1 μm, lateral junction; 0 μm, basal plane. The yellow box is shown as 2-color overlay XY slices in Fig. 2 and as single channels here (Scale bar, 1 μm). Dashed lines show the sectioning planes of XZ and YZ in Fig. 2. The cyan box is shown in **D**. (**D**) Raw images. Membrane concentrates to actin protrusions (arrows) but not the voids between protrusions (arrowheads). The plots show the density of membrane and the actin:membrane ratio. Scale bar, 5 μm (**A, C**), 1 μm (**B, D**).

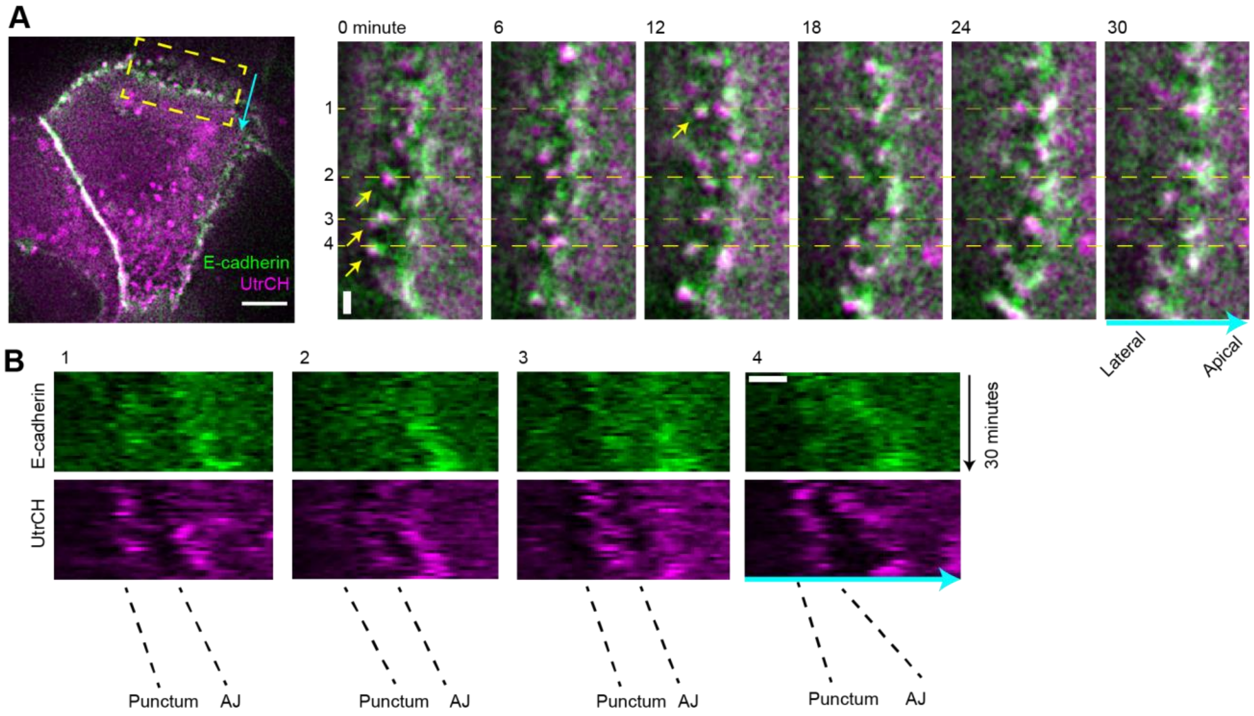


**Fig. S5. Membrane puncta are stable and do not move within the lateral membrane.**

(**A**) Left, a tilted region of lateral membrane (yellow box) in a cell coexpressing markers for actin (magenta) and E-cadherin (green). (Scale bar, 5 µm.) Right, the yellow box tracked in the time series. Membrane puncta (yellow arrows) colocalize with actin. These puncta do not diffuse within the lateral membrane in a basal-to-apical direction (cyan arrow and yellow dashed lines) in MDCK cells confirming the previous report (*23*). (Scale bar, 1 µm.) (**B**) Linescan of positions 1–4 (yellow dashed lines) are aligned frame by frame to generate kymographs shown here. Black dashed lines show the position of the punctum and apical junction. The distance between puncta and apical junction are relative constant over 30 minutes. Scale bar, 1 µm. See Movie S4.

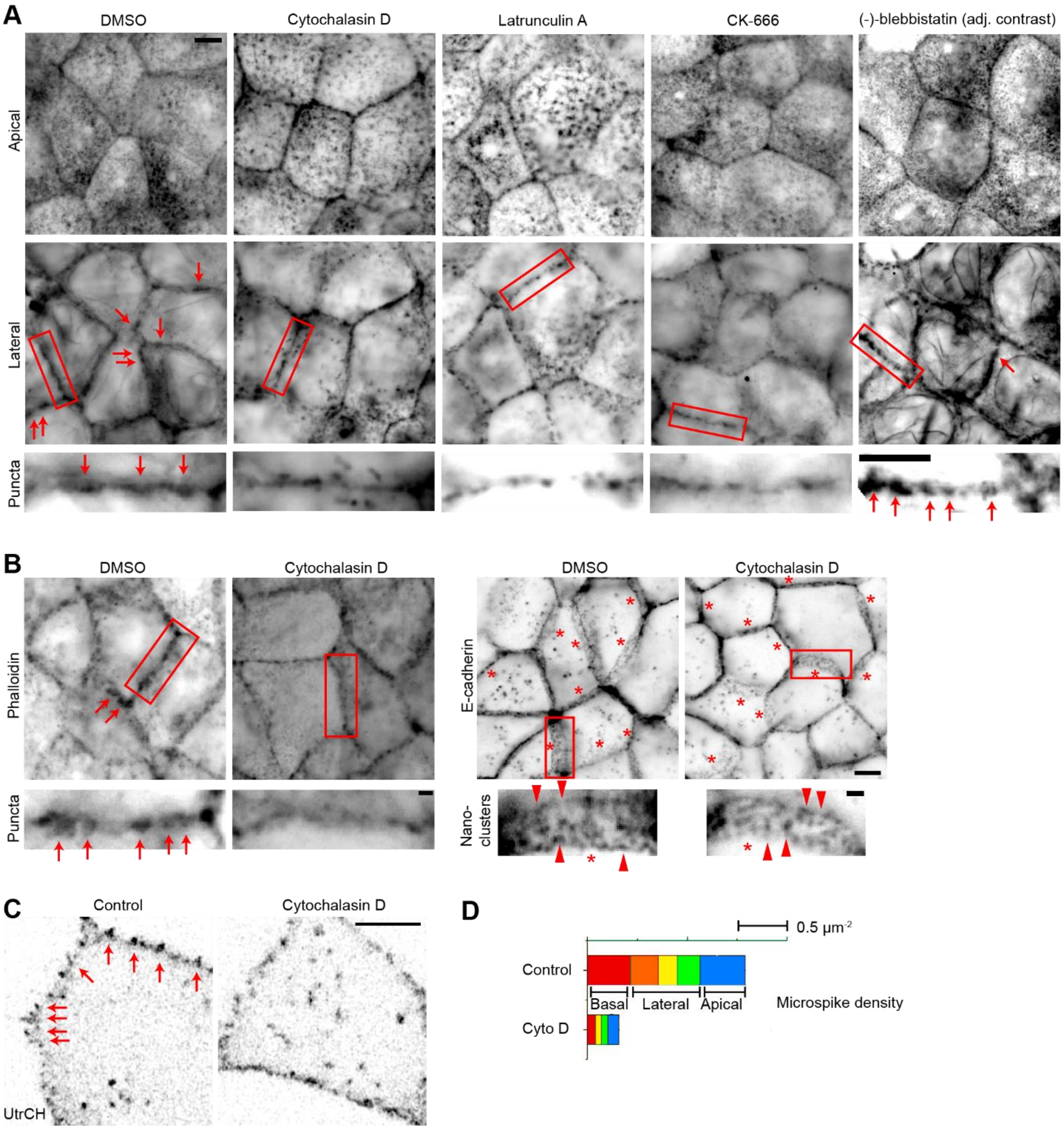


**Fig. S6. Actin puncta but not cadherin nanoclusters are sensitive to inhibition of actin polymerization.**

(**A**) Cytochalasin D (actin filament plus end capper at 200 nM, 0.5 h), latrunculin A (actin monomer sequester, 100 nM, 1 h), and CK-666 (Arp2/3 inhibitor, 100 µM, 1 h) remove the mesoscale actin puncta (arrows) from the lateral cortex, whereas (–)-blebbistatin (myosin II ATPase inhibitor, 50 µM, 1 h) does not. Dimethylsulfoxide, vehicle. The 4% formaldehyde used here is optimal for fixing mesoscale actin puncta. See figs. S9, S10, and S15 for validation of these drugs. Scale bar, 5 µm. (**B**) Cytochalasin D (25 nM, 0.5 h) removes the mesoscale actin puncta (arrows) but not the nanoscale cadherin clusters (arrowheads, DECMA-1 antibody) from the lateral surface. The apical side of the tilted lateral membranes is labeled by asterisks. See fig. S10 for validation of the drug treatment. Scale bar, 5 µm (top) and 1 µm (bottom). (**C**) Cytochalasin D (200 nM, 20 minutes) removes lateral microspikes (arrows) in GFP-UtrCH expressing cells that are surrounded by unlabeled cells. Scale bar, 5 µm. (**D**) The microspike density across the lateral membrane, which is normalized to membrane area in each z segment 1 µm deep (color blocks). Comparing all microspikes between control and cyto D, $P < 10^{-3}$; comparing apical junctional microspikes, $P < 0.004$; comparing lateral junctional microspikes, $P < 10^{-3}$, 2-sided t-test. $N = 10$ control and 8 cyto D treated junctions. Even in control, microspike density on lateral membranes is at least ~20 times lower than that is needed to densely pack the membrane surface with microspikes (neck diameter ~0.5 µm, see Methods for calculation).

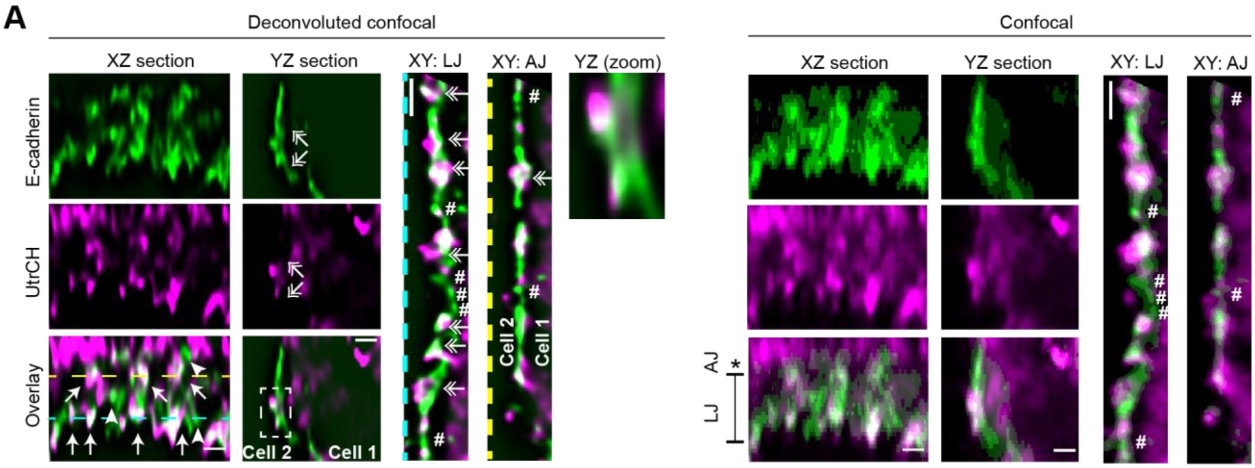


**Fig. S7. Lateral microspikes point outward and are tracked by E-cadherin.**

(**A**) Vertical and horizontal sections of a cell co-expressing markers for actin (magenta) and E-cadherin (green). This cell (cell 1) is surrounded by non-labeled cells including cell 2. Note all microspikes point to the left (outside of cell 1's cell body) in YZ and XY sections (double arrows). Microspikes may appear as mesoscale puncta before deconvolution. Dashed lines mark the sectioning planes. Arrows and arrowheads show E-cadherin puncta that are colocalized with actin or not, respectively. Asterisk and vertical dumb bell bar respectively mark the range of apical junction and lateral junction that are grazed by the XZ plane in which the XZ section is viewed en face. #, nanoscale cadherin clusters (monomers or oligomers) that are seen in both raw and deconvoluted images but are not always associated with actin. Scale bars, 1 µm.

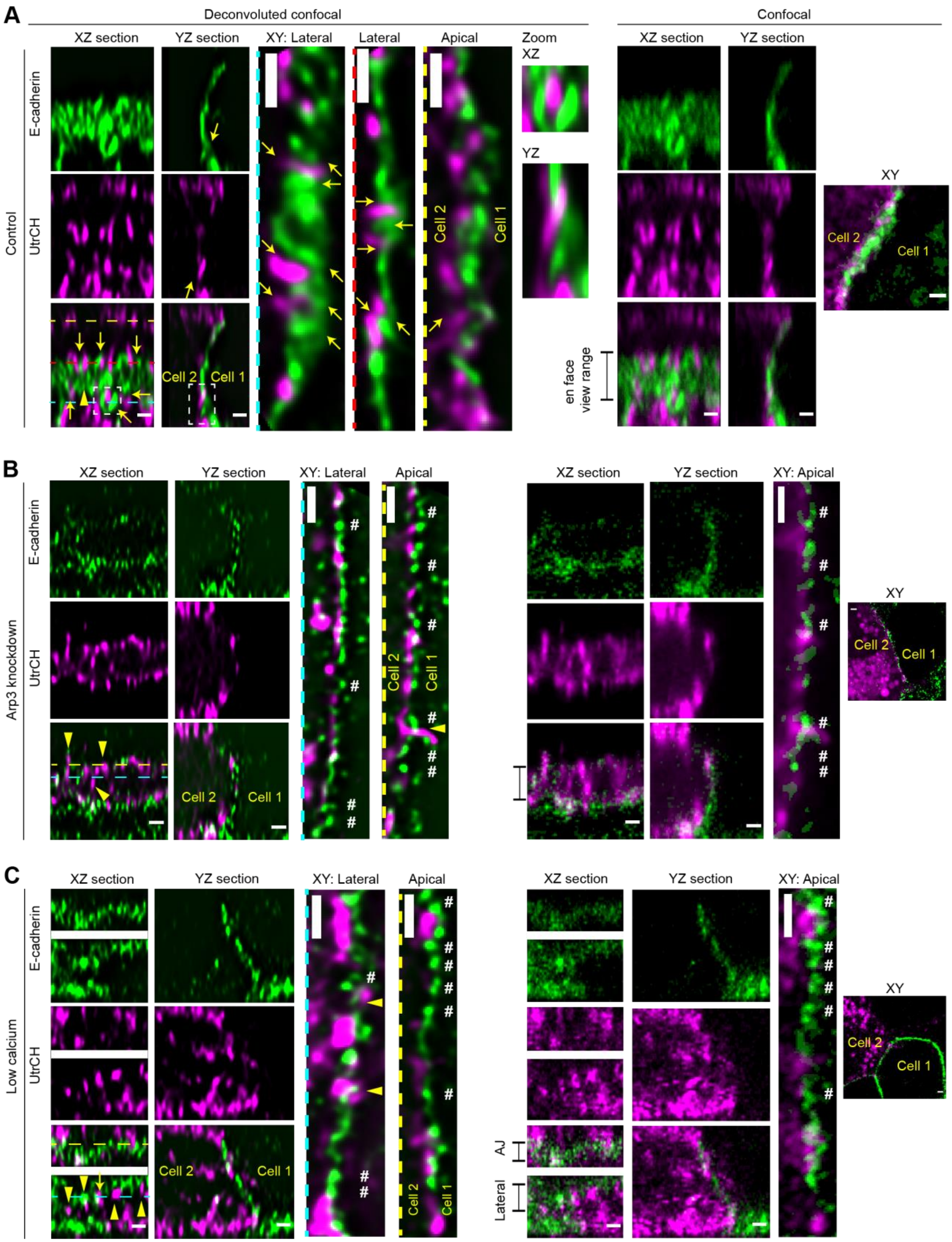
A
Deconvoluted confocal
Confocal
XZ section
YZ section
XY: Lateral
Lateral
Apical
Zoom
XZ
YZ
XY
Control
E-cadherin
UtrCH
Cell 2
Cell 1
en face view range
B
XY: Apical
Arp3 knockdown
C
Low calcium
AJ
Lateral

Fig. S8

**Fig. S8. Interdigitation between microspikes from neighboring 2 cells.**

(**A**) Vertical and horizontal sections of 2 cells expressing markers for E-cadherin (green, cell 1) and actin (magenta, cell 2) in confluent cell sheets. Despite microspikes are at least ~20 times lower than that is needed to densely pack the membrane surface (fig. S6), they cluster with one another suggesting they are kept together by some mechanism. Dashed lines mark the sectioning planes. Arrows and arrowheads show the mesoscale puncta that are clustered with mesoscale puncta on another cell or not, respectively. Asterisk and vertical dumb bell bar respectively mark the range of apical junction and lateral membrane that are grazed by the XZ plane in which the XZ section is viewed en face. Note the microspikes are short and side by side, unlike the contractile actin fibers that are much longer and connected head-to-head at a different type of cadherin mediated contacts referred to as “punctate” or “focal” adherens junctions (*24–30*). (**B**) Same as in **A** but the cell sheet expresses a small interference RNA against Arp3. (**C**) Same as in **A** but treated in low calcium media (90 µM, 1 d). #, the presence of nanoscale cadherin clusters, which is unlike the loss of mesoscale cadherin puncta (interdigitated microspikes). All scale bars, 1 µm.

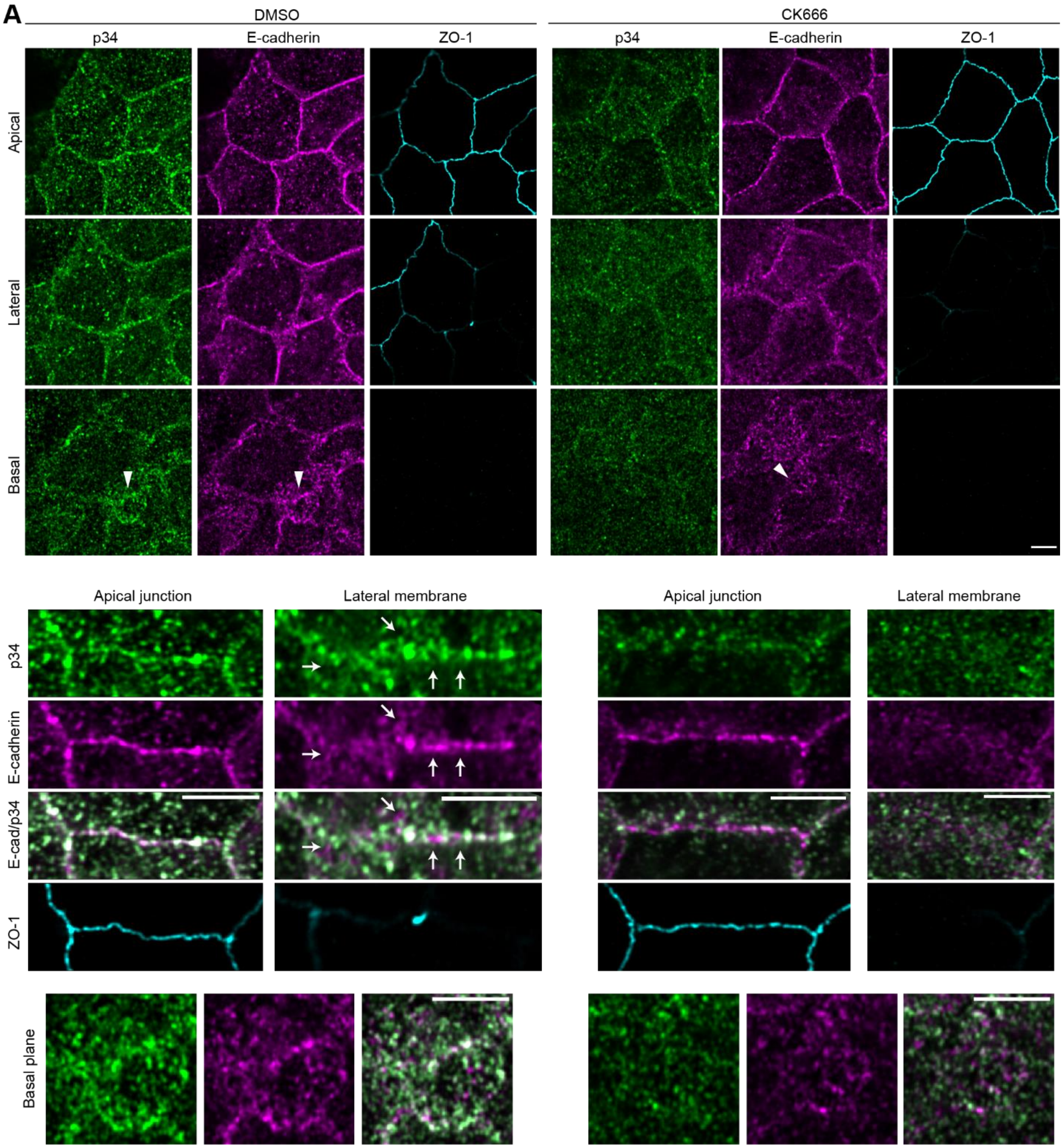


**Fig. S9. Arp2/3 is in the lateral cortex.**

(**A**) Arp2/3 subunit p34 localization in cell sheets treated with vehicle (dimethylsulfoxide) or Arp2/3 inhibitor CK-666 (100 µM, 1 h). The p34 antibody requires methanol fixation. This experiment also validates the specificity of the antibody and the effects of the drug. Note that p34 largely colocalizes with E-cadherin (rr1 antibody) at apical and lateral junctions with the exceptions marked by arrows. Note the E-cadherin nanoclusters but not p34 persisted after the drug treatment. Arrowheads, basal cryptic lamellipodia. Scale bar, 5 µm.

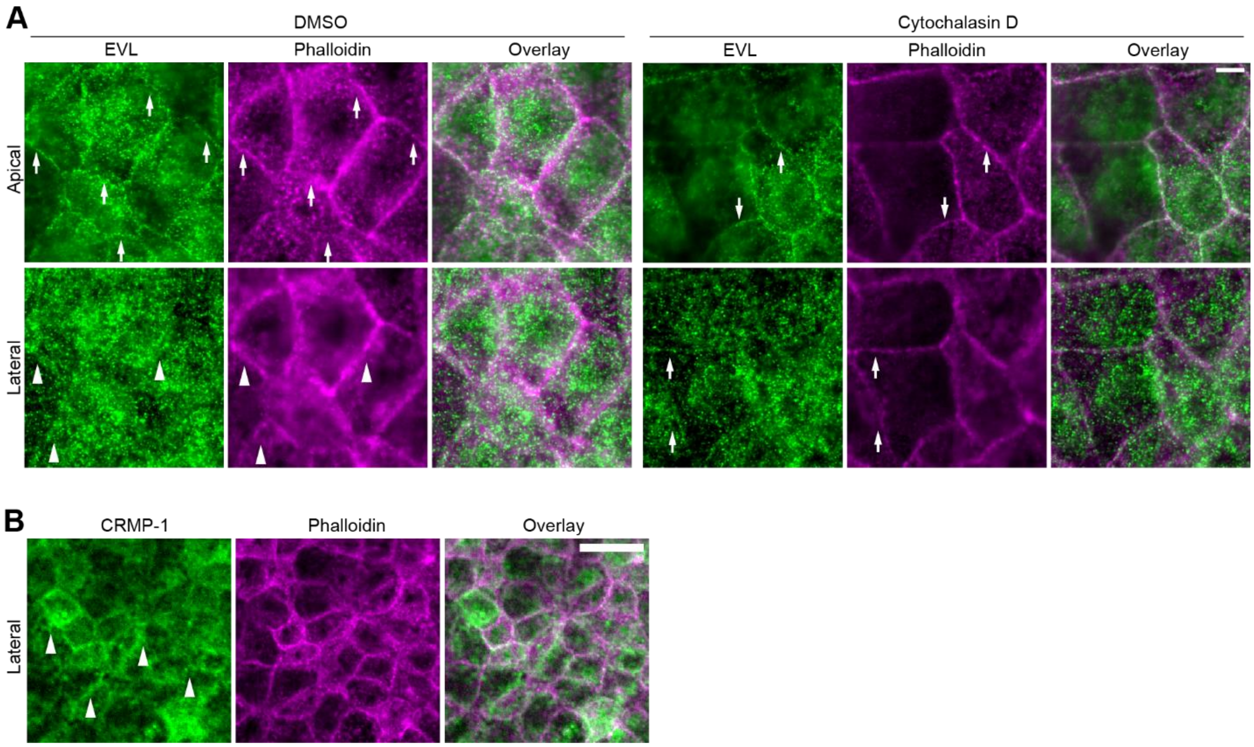


**Fig. S10. EVL and CRMP-1 are in the lateral cortex.**

(**A and B**) Endogenous EVL and CRMP-1 staining by simultaneous permeabilization/fixation. (**A**) Wild type cells treated with dimethylsulfoxide (vehicle) or cytochalasin D (25 nM). EVL and phalloidin staining at apical junctions (arrows) and lateral junctions (arrowheads) are decreased by low dose cyto D. Note the left part of the cyto D images are out of focus in the top panel; thus their apical junctions appear in the lower panel. This experiment also validates the effects of cyto D. Scale bar, 5 μm. (**B**) Arrowheads, lateral junctions. Scale bar, 25 μm.

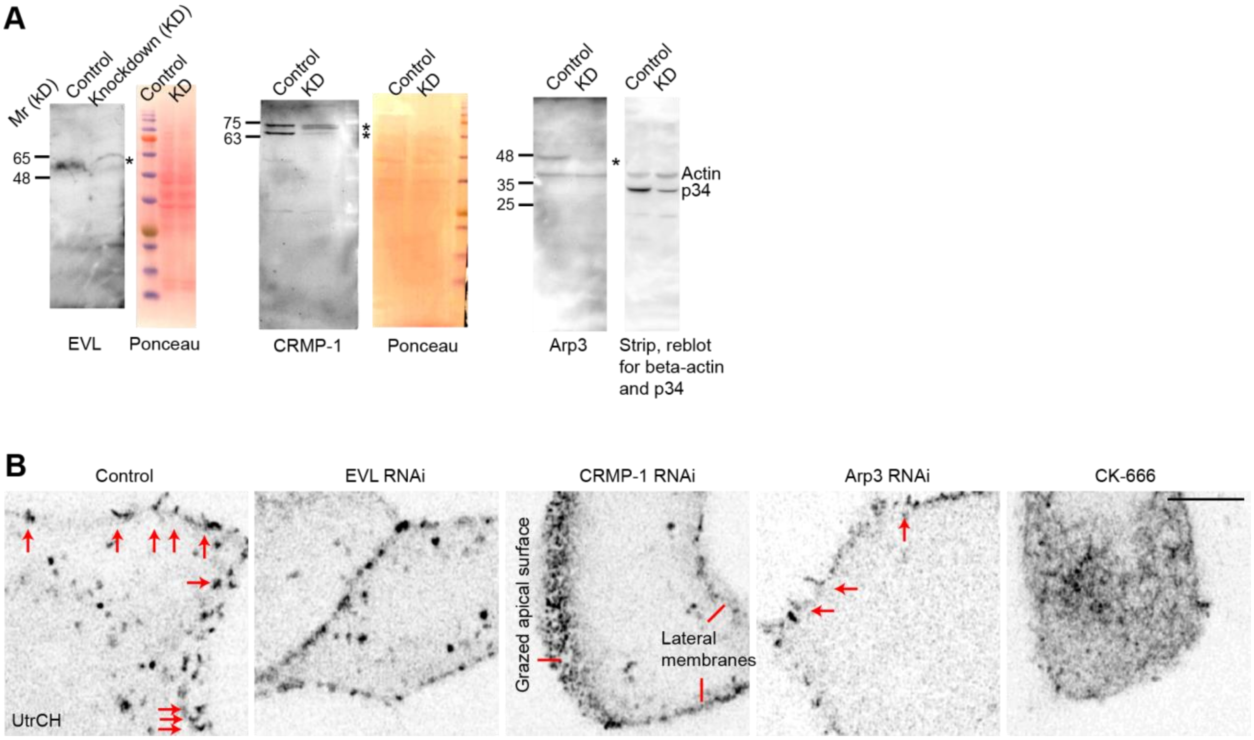


**Fig. S11. Actin polymerization factors regulate lateral microspikes.**

(**A**) Validation of knockdown. Equal total protein is loaded. The lower, unspecific band in Arp3 blot is likely actin, which is structurally like Arp3. p34 is another subunit of Arp2/3 complex. (**B**) Microspikes (arrows) in cells expressing mChe-UtrCH that are surrounded by non-expressing cells. Cropped images are shown in Fig. 4. See fig. S9 for validation of CK-666 treatment (100 µM, 0.5 h). Scale bar, 5 µm.

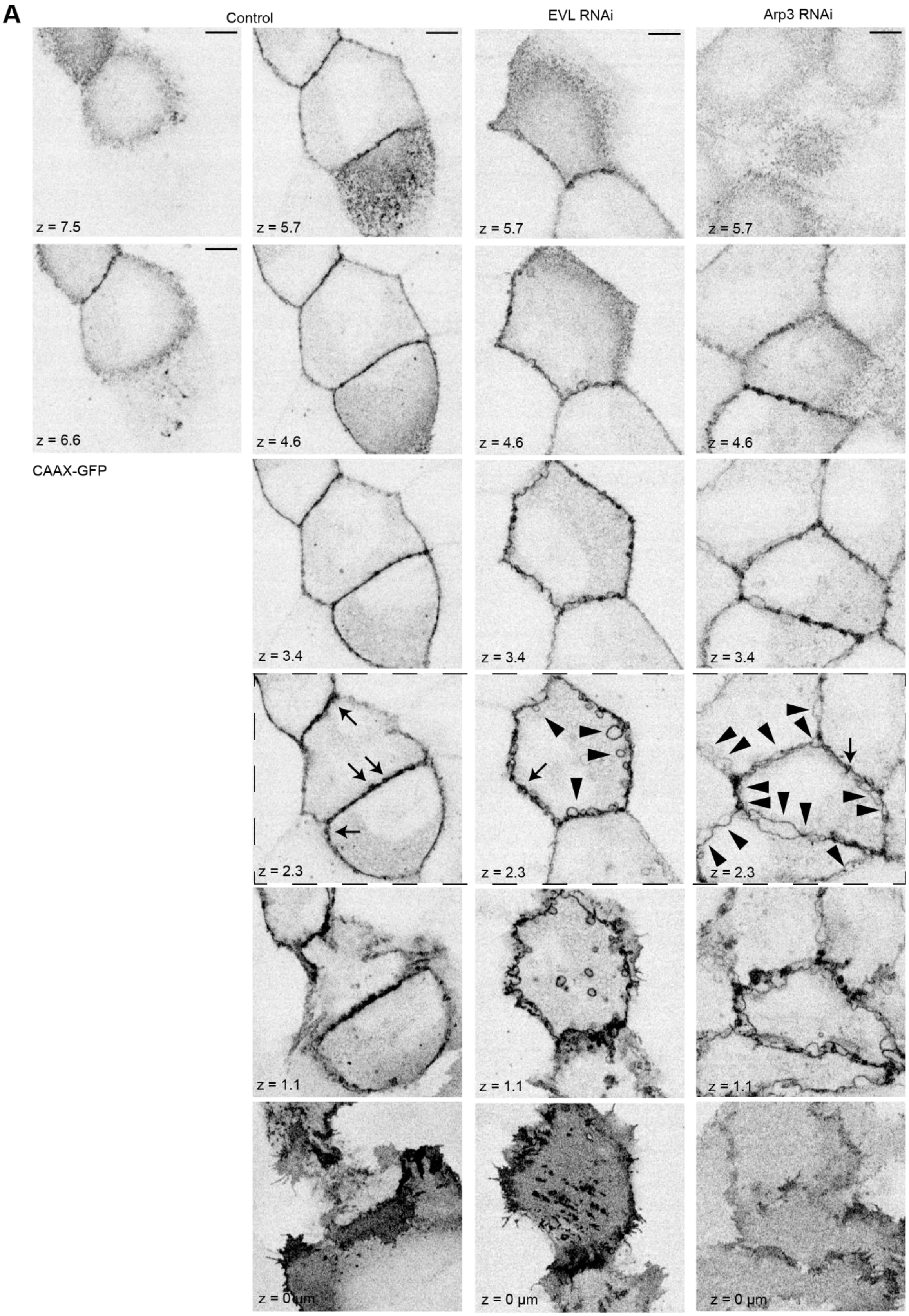
A
Control
EVL RNAi
Arp3 RNAi
z = 7.5
z = 5.7
z = 5.7
z = 5.7
z = 6.6
z = 4.6
z = 4.6
z = 4.6
CAAX-GFP
z = 3.4
z = 3.4
z = 3.4
z = 2.3
z = 2.3
z = 2.3
z = 1.1
z = 1.1
z = 1.1
z = 0 µm
z = 0 µm
z = 0 µm

**Fig. S12**

**Fig. S12. Membrane puncta and hernias in cell sheets lacking actin polymerization.**
(**A**) Z slices of plasma membranes in live cell sheets. Arrows, puncta. Arrowheads, hernias. Dashed line boxed regions are shown in Fig. 4. Scale bar, 5 μm.

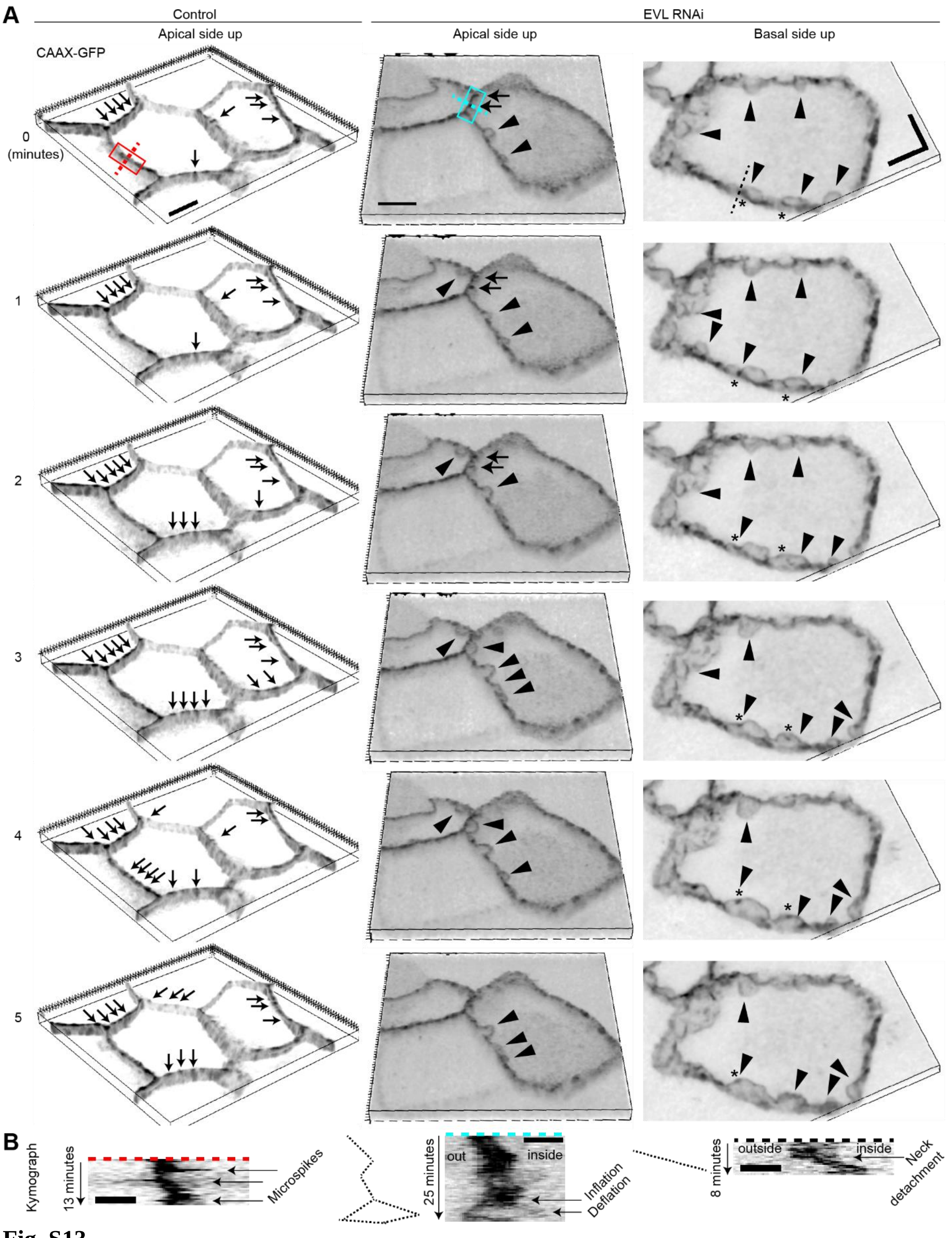
A
Control
EVL RNAi
Apical side up
Apical side up
Basal side up
CAAX-GFP
0
(minutes)
1
2
3
4
5
B
Kymograph
13 minutes
Microspikes
25 minutes
out
inside
Inflation
Deflation
8 minutes
outside
inside
Neck
detachment

**Fig. S13**

**Fig. S13. Membrane puncta are unstable in EVL knockdown cells and inflate into hernias.**

(**A**) Three-dimensional reconstruction of puncta (arrows) and hernias (arrowheads) on the plasma membranes. Right, 2 neighboring hernias merge and adopt a smooth surface suggesting the hernias are likely inflated by extracellular pressure (*6*); dense patches (asterisks) at the neck of the 2 hernias sometimes remain after herniation. Scale bar, 5 µm. Red box is shown in Fig. 2. Cyan box is shown in Fig. 4. (**B**) Kymograph of the dashed lines in **A** shows microspikes protruding in both directions from a punctum (left), a punctum inflating into a hernia (middle), and 2 hernias merge after their neck detach (right). Scale bar, 1 µm. See Movies S3, S6–S9.

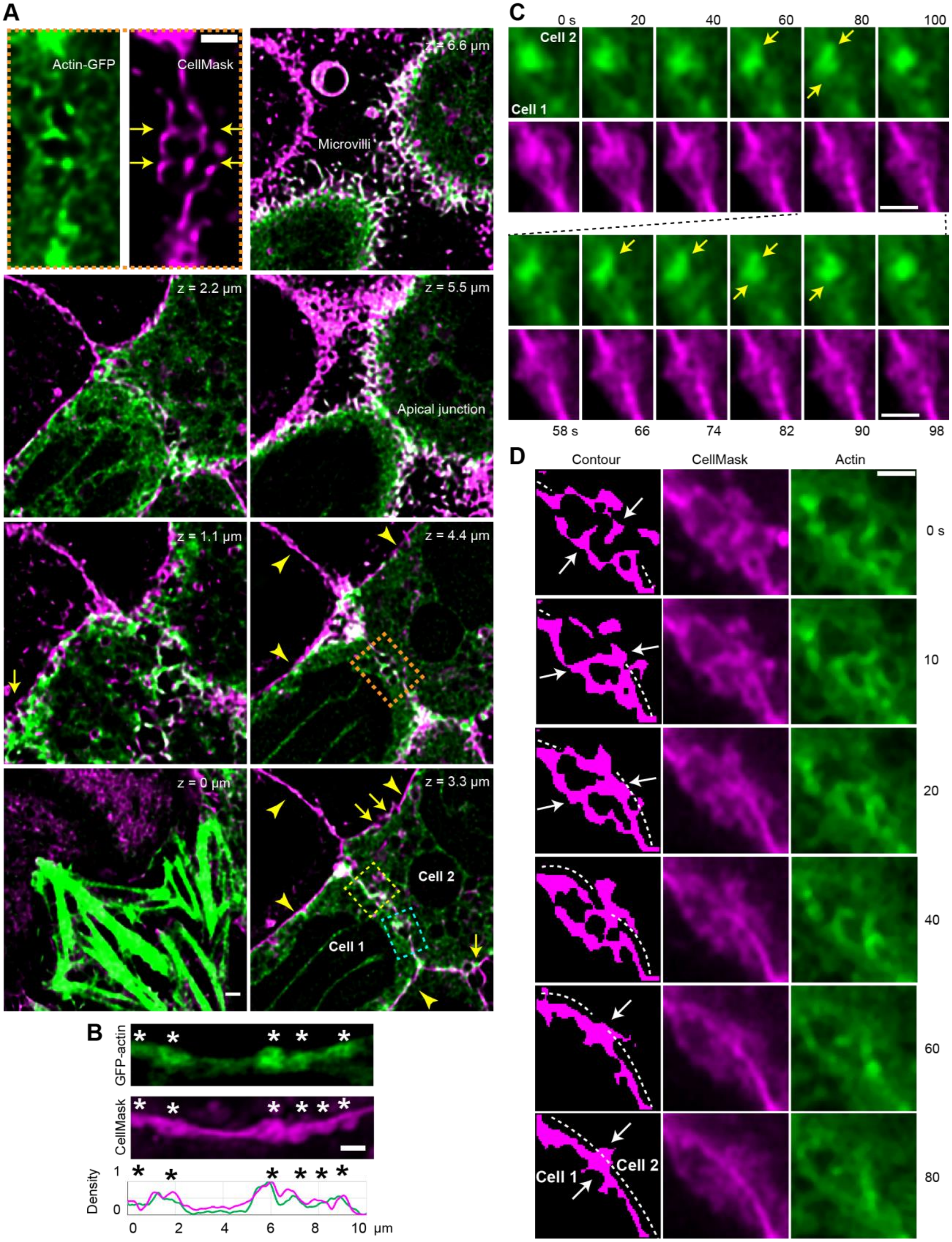
A
Actin-GFP
CellMask
z = 6.6 µm
Microvilli
z = 2.2 µm
z = 5.5 µm
Apical junction
z = 1.1 µm
z = 4.4 µm
z = 0 µm
z = 3.3 µm
Cell 2
Cell 1
B
GFP-actin
CellMask
Density
0 2 4 6 8 10 µm
C
0 s 20 40 60 80 100
58 s 66 74 82 90 98
D
Contour
CellMask
Actin
0 s
10
20
40
60
80

Fig. S14

**Fig. S14. Actin microspikes and the remodeling of convoluted membranes.**

(**A**) Control cell sheet expressing actin-GFP (green) is stained and imaged alive with CellMask (magenta). The junction of interest is of homogenous labeling. Arrows show engaged microspikes from 2 cells at detached membranes. Arrowheads show normal, adherent lateral membranes between 2 cells. Yellow and cyan boxes are shown in panel **C** and **D**. (**B**) Co-distribution of the 2 markers in lateral puncta (asterisks). (**C**) Microspikes (arrows) protrude in the antiparallel directions at a region of lateral membranes that started out as detached, convoluted but undergoes remodeling and flattening. (**D**) Microspikes from 2 cells search (0 s), capture (10 s), and stay engaged with each other (20 s and beyond), which is associated zippering of the herniated membranes. The contour is generated from the membrane channel which shows the adherent region of 2 lateral membranes traced with dashed lines. All scale bars, 1 µm. See Movie S10.

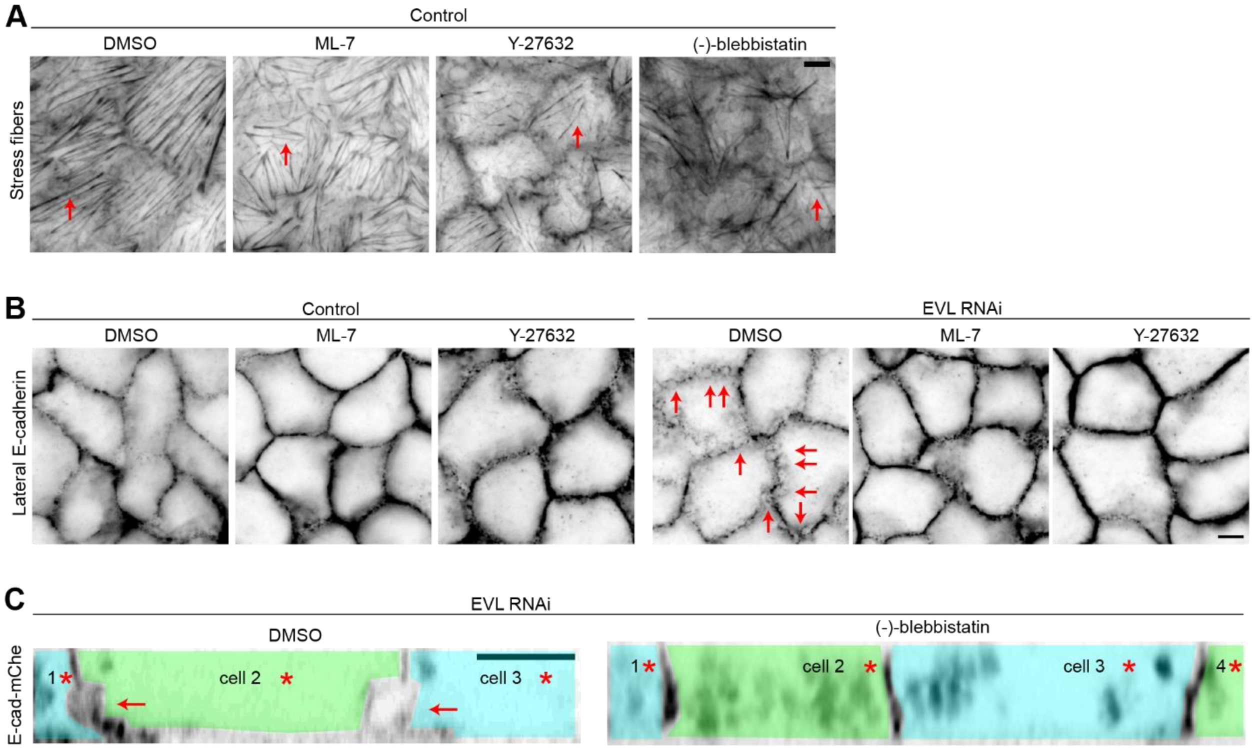


**Fig. S15. Myosin II contractility does not maintain cell-cell adhesion.**

(**A**) Validation of drug treatment. Basal stress fibers (arrows) become thin and disappear after treated by ML-7 (Myosin Light Chain Kinase/MLCK inhibitor, 25 µM, 0.5 h), Y-27632 (Rho-associated Kinase/ROCK inhibitor, 25 µM, 0.5 h) or (–)-blebbistatin (myosin II ATPase inhibitor, 50 µM, 0.5 h). Dimethylsulfoxide, vehicle. (**B**) Lateral z slices of E-cadherin staining. Methanol fixation. Inhibiting contractility does not disrupt cell-cell adhesion in wild type; it rescues herniation (arrows) in EVL knockdown cells. (**C**) Live cell sheet vertical sections where cells are shaded with pseudo colors. Asterisks, marker expressing cells. Inhibiting contractility rescues herniation (arrows) in EVL knockdown cells. Scale bar, 5 µm.

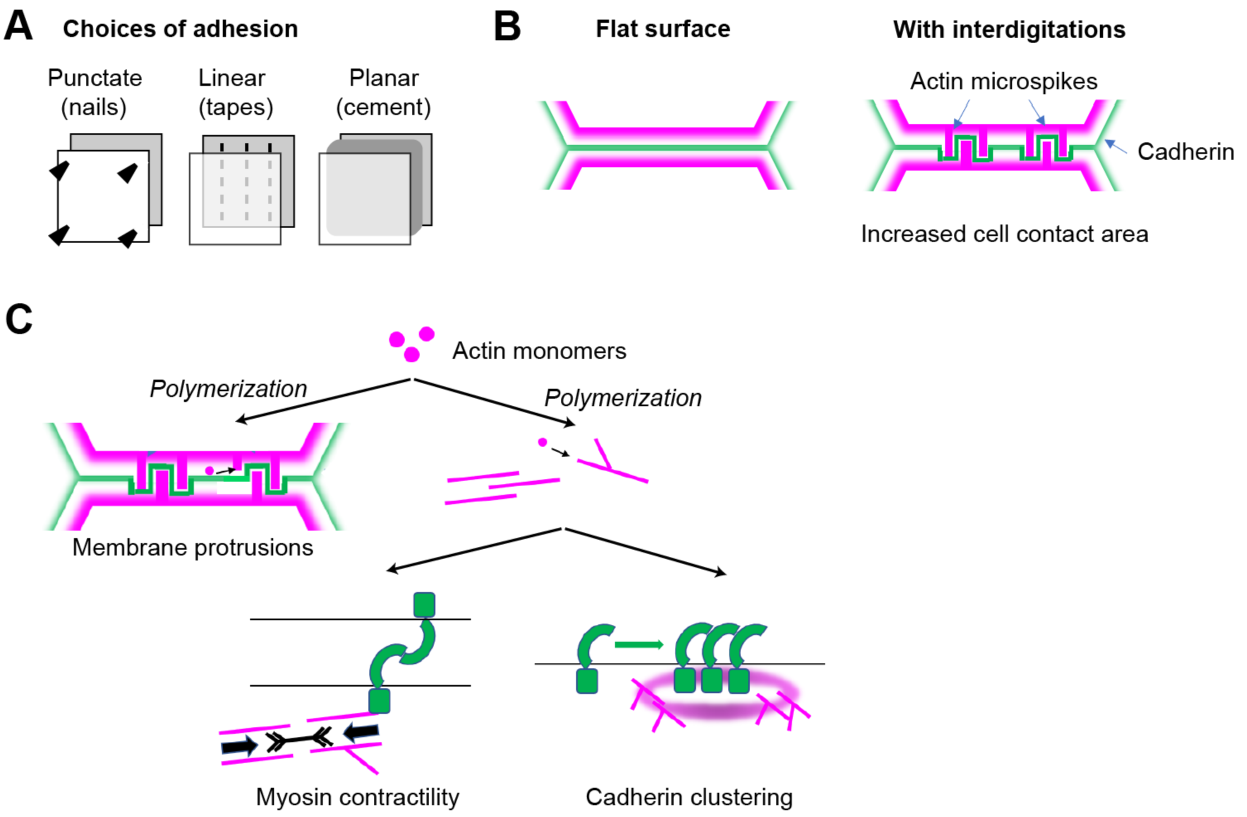


**Fig. S16. The actin protrusion model of cadherin adhesion maintenance.**

(**A**) Three choices to keep the expansive lateral plasma membrane surfaces attached: the punctate (e.g. microspikes and cellular interdigitations), linear (e.g. cadherin dependent adherens and desmosomal junctions), and planar (e.g. plant cell walls) forms of contacts. (**B**) Interdigitated actin microspikes emerging from actin cortex (magenta) greatly increase the cell contact area for cadherin (green) mediated adhesion. (**C**) The actin protrusion model adds less entities than the contractility model and the clustering model to explain the continuous actin polymerization at mature junctions, thus it would better survive Occam's razor. Actin polymerization can directly drive membrane protrusions to keep two cells' cadherins close in favor of homophilic bonds; although filaments generated by polymerization may contribute to contractility or clustering, these functions are not the immediate consequences of actin polymerization per se. Further, the actin to drive membrane protrusions must be a dynamic population, agreeing with experimental observations; yet the actin filaments as the tracks for myosin or the tethers and fences for cadherins do not necessarily need to be dynamic.

**Table S1. Perturbations of lateral microspikes and hernias.**

| | Microspikes | Hernias |
|---|---|---|
| EVL knockdown | – | + |
| CRMP-1 knockdown | – | + |
| Arp3 knockdown | – | + |
| CK-666 | – | ND |
| (–)-Blebbistatin | + | – |
| ML-7 | + | (note 1) |
| Y-27632 | ND | (note 1) |
| Low calcium | ± (note 2) | + |
| Cytochalasin D | – | ND |
| Latrunculin A | ND (note 3) | + |

+, increase; –, decrease; ±, no effect, compared to control; ND, not determined.
Note 1: these treatments rescue hernias in EVL and Arp3 knockdown cells.
Note 2: this treatment decreases microspike engagement.
Note 3: this treatment decreases actin puncta.

**Movie S1.**

Fluorescence recovery after photobleaching of E-cadherin-mCherry in a flat region (left) and a punctum (right). Frame interval, 0.4 s.

**Movie S2.**

Fluorescence recovery after photobleaching of CAAX-GFP in a punctum. Frame interval, 0.166 s.

**Movie S3.**

Membrane puncta and protrusions labeled by CAAX-GFP. Frame interval is 30 s, shown as min:sec.

**Movie S4.**

The apical junction and a tilted part of lateral membrane in a cell co-expressing E-cadherin-GFP (green) and mCherry-UtrCH (magenta). Neighboring cells are dark. Frame interval is 60 s, shown as min:sec.

**Movie S5.**

The left and right cell expresses E-cadherin-GFP (green) and mCherry-UtrCH (magenta), respectively. Frame interval, 1 s.

**Movie S6.**

3-dimensional reconstruction of lateral membranes labeled by CAAX-GFP. Control MDCK cell sheet viewed from apical side. Frame interval, 30 s. Compare with Movies S7 and S8.

**Movie S7.**

3-dimensional reconstruction of lateral membranes labeled by CAAX-GFP. EVL knockdown cell sheet viewed from apical side. Frame interval, 30 s. Compare with Movie S6.

**Movie S8.**

3-dimensional reconstruction of lateral membranes labeled by CAAX-GFP. EVL knockdown cell sheet viewed from basal side. Frame interval, 30 s. Compare with Movie S6.

**Movie S9.**

A membrane punctum labeled by CAAX-GFP bursts into a hernia. EVL knockdown cell sheet. Frame interval is 30 s, shown as min:sec.

**Movie S10.**

GFP-β-actin (green) and a lipophilic dye CellMask (magenta) showing microspikes on the 2 cells' herniated membranes reengage each other. Frame interval, 1 s.
